# A context-free grammar for Caenorhabditis elegans behavior

**DOI:** 10.1101/708891

**Authors:** Saurabh Gupta, Alex Gomez-Marin

## Abstract

Hierarchy is a candidate organizing principle of ethology, where actions grouped into higher order chunks combine in specific ways to generate adaptive behavior. However, demonstrations of hierarchical organization in behavior have been scarce. Moreover, it remains unclear how such underlying organization allows for behavioral flexibility. Here we uncover the hierarchical and flexible nature of *Caenorhabditis elegans* behavior. By describing worm locomotion as a sequence of discrete postural templates, we identified chunks containing mutually substitutable postures along the dynamics. We then elucidated the rules governing their interactions. We found that stereotypical roaming can be described by a specific sequence of postural chunks, which exhibit flexibility at the lowest postural level. The same chunks get combined differently to produce dwelling, capturing non-stereotypical actions across timescales. We show that worm foraging is organized hierarchically —a feature not explainable via Markovian dynamics—, and derive a context-free grammar governing its behavior —which is different than a regular grammar, or a hidden Markov chain. In sum, in making the analogy with human language concrete (but not literal) our work demonstrates, in line with the foundational insights of classical ethologists, that spontaneous behavior is orderly flexible. Once more, investigating the humble nematode suggests that everything human has its roots in lower animal behavior.

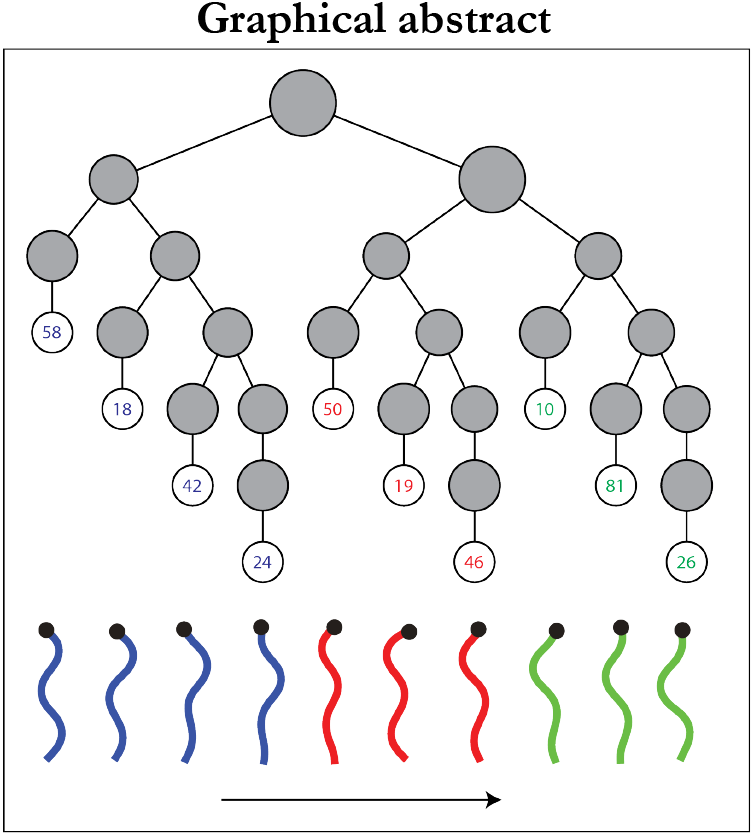
Graphical abstract.

## 1. Introduction

In order to achieve a variety of goals, we and other animals express complex behaviors in terms of action sequences. Current methods allow for automated high-resolution behavioral measurements, generating big datasets that in turn pose new challenges and opportunities for behavioral analyses (**Perona & Anderson, 2014; Gomez-Marin et al., 2014**). Understanding the organizational principles underlying the generation of innate or learned actions can then reveal computational primitives ascribable to their genetic and neural circuits underpinnings (**Gallistel, 1980; Gallistel, 1981**). By thoroughly working out algorithmic explanations of behavior, the quest for its mechanistic bases shall become more meaningful and rewarding (**Karkauer et al., 2017**).

Animal behavior is not only variable, but saliently flexible. The number of neurons, muscles and joints are orders of magnitude greater than required to make desired movements, posing the degree of freedom problem for motor systems (**Bernstein, 1967**). Such redundancy is a significant challenge towards understanding behavior, since different motor signals can generate the same behavior (degeneracy), while similar motor signals can generate completely different behaviors based on the context in which they appear (reusability) (**Tononi et al., 1999; Sporns & Edelman, 1993; Bernstein, 1967**). It is interesting to note that degeneracy and reusability can lead to flexibility, even solely at the level of behavior. For instance, we can brush our teeth either by moving the head up-and-down or by keeping it still while moving the hand up-and-down. And we can move the hand up-and-down while brushing our teeth as well as while cleaning a window. Thus any principle of behavioral organization must also account for this “either/or” feature of behavior where distinct sequences of actions can be substituted with each other to generate flexible behavior.

Hierarchical organization has been postulated as a general principle of behavior (**Tinbergen, 1950, 1951; Simon, 1962; Dawkins, 1976; Lashley, 1951**) that can also tame the redundancy problem while generating flexible behaviors (**Dawkins, 1976; Dawkins & Dawkins, 1973**). In the words of Herbert Simon, “*hierarchy, I shall argue, is one of the central structural schemes that the architect of complexity uses”* (**Simon, 1962; Rosenbaum et al., 2007**). In turn, Karl Lashley (**Lashley, 1951**) had hypothesized that behavioral sequences are generated by a hierarchical organizational schematic, whereby low level behavioral descriptions (like words) are grouped together into higher order behavioral descriptions, i.e. chunks (like phrases), that further get grouped into still higher order “chunks of chunks” (sentences).

This is in sharp contrast with a linear chain view of behavioral sequence generation, where the preceding action in a sequence triggers the next. Such is the Markovian assumption that pervades a great family of models. For instance, the ambiguity in the phrase *‘white taxi driver’* can be easily expressed in a hierarchical schematic but it is impossible to be expressed in a linear system such as one consisting of transition probabilities between words (Markov model) or even between categories of words (akin to a hidden Markov model) (**Dehaene et al., 2015**). As depicted in **Figure 1**, *‘White taxi driver’* is a chunk that in turn consists of two potential sub-chunks, namely, *‘taxi driver’* and *‘white taxi’.*

**Figure 1.**
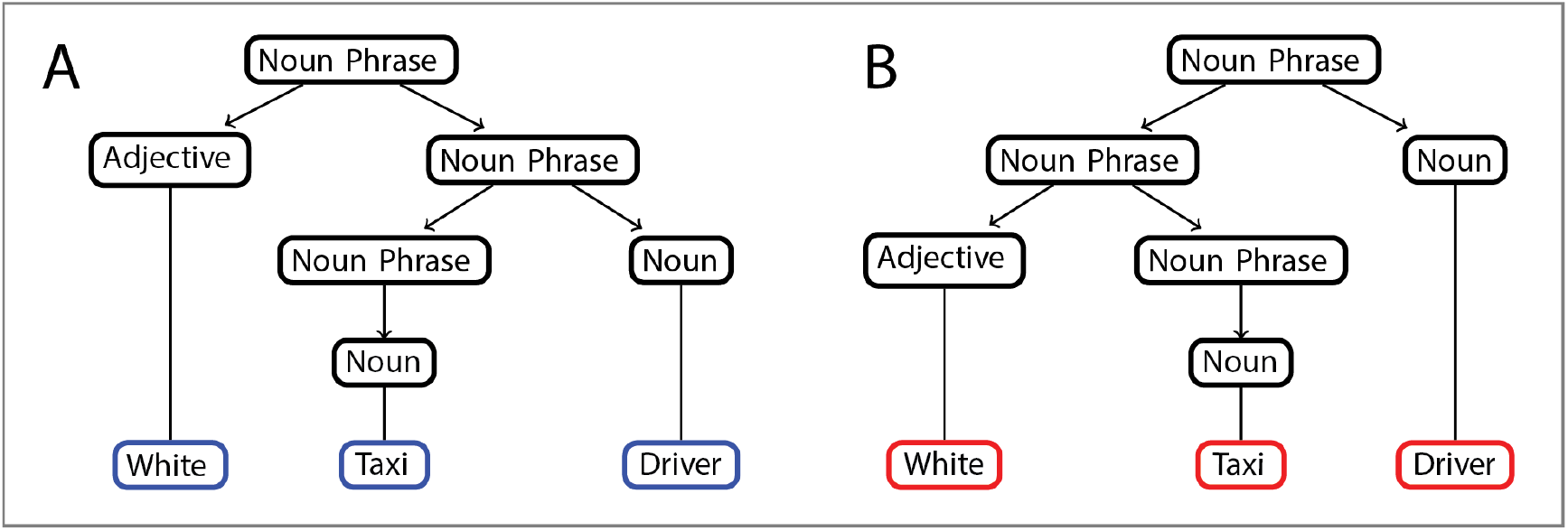
Flat structures fail to account for sequence production, syntax, and meaning. The ambiguity in the sentence “*White taxi driver*” cannot be accounted using flat sequential structures. In contrast, nested rules (in black) can reflect compositionality in hierarchical systems: the rules of grammar lead to an interpretation where the taxi driver is a white person (**A**); in a different interpretation of the same phrase, a driver drives a white taxi (**B**).

Moreover, the process of generating higher order chunks from sub-chunks follows a set of rules. For example, the sub-chunk *‘taxi driver’* when preceded by an adjective, *‘white’*, gives one meaning to the phrase ‘*white taxi driver’* and when the sub-chunk *‘white taxi’* is succeeded by the noun *‘driver’*, another meaning arises. This highlights the compositional property of hierarchical systems where higher order chunks are defined in terms of simpler constituent components according to a set of production rules (such as “Noun Phrase → Adjective (Noun Phrase)” and “Noun Phrase → (Noun Phrase) Noun”). Again, compositionality endows flexibility to hierarchical systems by allowing the re-use of stable sub-chunks and chunks according to different rules. For example, the sub-chunk ‘*taxi driver’* above can be re-used to generate two different sentences: (1) The license of the ‘*taxi driver’* was confiscated, and (2) The ‘*taxi driver’* hit the person on the road and fled from the scene.

However, it remains unclear to what extent the richness of animal behavior can be captured by such a compositional hierarchy and whether that organization can yield deep insights—mechanistic, and also algorithmic— into ecologically relevant behaviors. Many of the ecologically important and fitness determining decisions that all animals make are related to foraging, which is one of the most critical problems faced by all animals (**Mobbs et al., 2018**). Modulating their foraging behavior according to internal and external conditions is critical for the survival and future reproduction of animals (**Cohen et al., 2009**). Thus, studying foraging behavior in the nematode worm *Caenorhabditis elegans* presents a promising direction to uncover the molecular and neural basis of this universal behavior, which could help decipher decision-making principles in humans as well.

The experimental tractability of *Caenorhabditis elegans* as a model organism holds considerable promise in elucidating molecular and neural mechanisms underlying locomotion. Indeed, receptors important for worm roaming behavior have been shown to have similarities with receptors that modulate feeding behavior in mammals (**Bendena et al., 2008; Cohen et al., 2009**). For example, the neuropeptide receptor *npr-9* has been shown to affect *Caenorhabditis elegans* behavior by inhibiting dwelling behavior, and is most similar to mammalian galanin receptors known to modulate feeding behavior in mammals. It is unclear if other neuropeptide receptors might also be involved in modulating worm foraging.

*Caenorhabditis elegans* foraging behavior is thought to be organized into two distinct states: the exploratory phase of roaming and the exploitative phase of dwelling (**Fujiwara et al., 2002; Ben Arous et al., 2009**). During roaming, the worm moves quickly across the bacterial lawn of food with low frequency of turns, whereas during dwelling, it moves rather slowly with frequent turns, thus confining itself to a very small region (**Flavell et al., 2013**). The same motor patterns (forward locomotion, reversals, turns) occur in roaming-specific or dwelling-specific combinations, giving both of these states their distinct characteristics (**Flavell et al., 2013**).

In search for the organization of worm behavior, iterative clustering of postural sequences during locomotion has been previously used in order to identify behavioral motifs and detect phenotypic differences between worms of different strains, or in different environments (**Brown et al., 2013; Gomez-Marin et al., 2016**). Such approaches can yield, by construction, hierarchical motifs but still leave open the question of hierarchy in worm behavior. In order to identify overrepresented and underrepresented sequence motifs when the environment is changed or when its nervous system is stimulated, an n-gram model has been used to fit worm behavior (**Schwarz et al., 2015**). However, these analysis frameworks, drawing on repeated action sequences, favor stereotypy at the expense of the flexibility commonly observed in *Caenorhabditis elegans* behavior (**Chao et al., 2004; Cheung et al., 2005; Chang et al., 2006**).

Markovian analyses are not circumscribed to worm behavior, but a commonly used approach to study the behavior of a variety of species, both within the classical ethology tradition and also in the latest sophisticated instantiations of computational ethology, such as in mice (**Wiltschko et al., 2015**). However, the behavior of organisms is a not single timescale phenomenon (**Berman, 2018**), as the multiplicity of timescales observed in fruit flies demonstrates (**Berman et al., 2014**). Indeed, the fruit fly has been a good organism model to investigate hierarchical organization in animal behavior, especially in the context of grooming (**Dawkins & Dawkins 1976; Seeds et al., 2014; Mueller et al., 2019**) as well as in courtship behavior (**McKellar et al., 2019**), and also with respect to its global postural behavior (**Berman et al., 2016**).

It is still unclear whether hierarchical organization —where primitive behavioral descriptions get grouped into chunks at a higher level of description along with the rules of interaction between those chunks (in a compositional hierarchy)— can explain *Caenorhabditis elegans* foraging behavior. We also lack understanding as to how the principle of compositional hierarchy can be tied to the flexible generation of *Caenorhabditis elegans* foraging behavior. These are the main goals and challenges of the present work.

Here, by using a publicly-available high-resolution database of *Caenorhabditis elegans* behavior (**Yemini et al., 2013**) and treating its movement as a discrete sequence of changes in its body posture (**Schwarz et al., 2015**), we show that worm locomotor sequences giving rise to foraging behavior are organized in accordance with a compositional hierarchy, including the non-stereotypical portions of its behavior. With the aim to capture degeneracy and reusability of behavioral elements, we use the idea of substitution (**Maurus & Pruscha, 1973; Dawkins, 1976**) to obtain chunks containing mutually substitutable worm postures. We then elucidate a grammar of worm roaming and dwelling states, outlining rules of interaction between such chunks. Next, we find that the stereotypical worm roaming behavior is captured by a specific grammatical rule involving specific chunks in a particular order. Even such stereotypical behavior is characterized by variability at the lowest level of postures. We also delineate grammatical rules that specify how the same chunks are re-used in different ways to produce relatively less stereotypical dwelling like behavior patterns. We show that the properties of the proposed grammatical rules are consistent with known experimental results about *Caenorhabditis elegans* foraging. Using the proposed grammar for worm foraging, we report hitherto uncharacterized role of neuropeptide receptors *npr-3* and *npr-10* in modulating *Caenorhabditis elegans* foraging behavior. In sum, a generative grammar for worm foraging demonstrates how flexible behavior can emerge from a compositional hierarchy.

## 2. Materials and methods

### Experimental data

We made use of the openly available *Caenorhabditis elegans* behavioral dataset that has been described previously (**Yemini et al., 2013**). Worms (N2, wild isolates, and several mutants) were picked to the centre of a patch of *Escherichia coli* OP50 on an agar plate, one at a time. The worms were allowed to habituate for 30 minutes before being tracked for a period of 15 minutes.

### Posture discretization

Following (**Schwarz et al., 2015**) we use a discrete representation of worm behavior where its locomotion is approximated by a sequence of discrete postures drawn from a finite number of postural templates. To do so, one first finds the angles of worm midlines at equally spaced points. Then, the skeleton angles in each frame are approximated to its closest matching postural template out of 90 templates that were derived from wild type N2-worms using k-means clustering. This results in the same template being fit to multiple consecutive frames, during which the animal might not be making huge changes to its posture. Thus, in order to disambiguate same postures repeated at different speeds, a simple non-uniform time warping procedure is used to swallow up repeats from the postural sequence. Thus the behavioral sequence {5, 5, 5, 4, 4, 3, 3, 2} is turned into {5, 4, 3, 2}. Note that the timing information of each posture is still conserved and can be used. In this way, worm foraging behavior is represented as a sequence of postures with each posture having timing information, denoting the amount of time the worm stayed in that posture before moving to the next one.

### Transition matrix

Considering all the 1287 individual “N2” worms on food, each of whose foraging behavior is described by a sequence of 90 postures, a first order 90×90 behavioral transition matrix B was created for all the worms pooled together. Each entry B_ij (i and j being subscripts in this notation) in the transition matrix denotes the number of times the worm made a transition from posture i to posture j across all the worms, pooled together.

### Mutual replaceability

As described in (**Maurus & Pruscha, 1973; Dawkins, 1976**), mutual repleaceability (MR) was applied to first-order Markov transition matrix to obtain chunks of mutually replaceable worm postures. To capture the various permutations in which postures can combine to produce sequences, MR seeks to cluster those behavior patterns together that do not necessarily occur close by in time, but whose transition relationships with members of other clusters are similar. Thus, these behavior patterns are mutually substitutable in those parts of the transition matrix, B, that do not involve their interactions with behavior patterns in their own cluster (**Dawkins, 1976**). For instance, two adjectives might usually not occur together in time in a sentence, but certain words in adjective cluster are mutually substitutable because they can be substituted in their interactions with other words in the noun cluster.

Starting with B, MR calculates index of mutual replaceability for each pair (i and j) of postures. Let the row in B, denoting the transition structure from posture i to all other postures be denoted as r_i and the row corresponding to posture j as r_j. Analogously, let the column in B corresponding to the transition structure from each of the 89 different postures to posture i be denoted as c_i and the column corresponding to posture j as c_j. For postures i and j, the Pearson correlation coefficient R_ij between r_i and r_j (excluding mutual interactions) and C_ij, the correlation coefficient between ci and cj (excluding mutual interactions between i and j) is first computed. The index of mutual replaceability, M_ij for i and j is finally computed as the mean of R_ij and C_ij.

The pair of postures, s and t, that has the maximum associated M_st value is then put in the same cluster and the transition matrix B is collapsed so as not to make any distinction between postures s and t by adding their corresponding entries. The same procedure is carried out on the reduced matrix, so that the next clustering might involve two different postures or one posture with an already made cluster in a previous iteration of the procedure. The procedure comes to a halt when only two entries remain in the ever reducing matrix. This process of iteratively forming clusters, gives rise to a dendrogram in which the portions that get merged higher up in the dendrogram are less substitutable than the pairs that get merged at a lower height in the tree. Cutting the tree formed at a certain height yields 10 visibly distinct clusters (b1, b2, b3, b4, r1, r2, r3, g1, g2, g3) which become the postural chunks and sub-chunks for our subsequent analyses.

### Silhouette values

The silhouette value of each posture p in a cluster is computed according to the equation Sp= (b_p−a_p)/max(a_p,b_p), where a_p denotes the average morphological distance between the p-th posture and all the other postures belonging to the same cluster as posture p, b_p denotes the minimum average morphological distance between posture p and all the other postures belonging to a different cluster than posture p, minimized over all the clusters. Euclidean distance d(p, q) used to compute the morphological distance between two postures p and q is defined as the square root of the sum of (p_i − q_i)^2 over all angles along each posture.

### Shuffling postures

In the control experiments with shuffled postures, each worm postural sequence is shuffled such that individual posture occurrence frequencies are maintained. For example: an original sequence {1, 2, 3, 1, 3, 4} might be shuffled as {3, 1, 2, 1, 4, 3}.

### Transition matrix for chunks obtained by MR

After applying MR on the postural sequence data of all the N2 worms combined, each of the 90 postures are assigned to one of the 10 clusters (b1, b2, b3, b4, r1, r2, r3, g1, g2, g3). Given a one-to-one mapping of each posture to one cluster, the postural sequences of the all the worms can be abstracted in terms of transitions between the 10 clusters. A transition from posture i to posture j is counted as a transition between clusters C_i and C_j, containing postures i and j respectively. In this way, a transition matrix between the 90 postures for the all the worms taken together can be transformed into a transition matrix between 10 clusters.

### Entropy

To analyze the transitions between postures and clusters, one can use entropy measures. Assuming that all the events are equi-probable, the uncertainty in predicting the next event (either posture or cluster) can be calculated in bits as H0=log(n), where the log is in base 2, and n is the number of values that can be taken by the variable under consideration. The reduced uncertainty that is afforded by the knowledge of individual event (posture or cluster) probabilities a priori is given by the usual H1=−sum[pi log(p_i)], where p_i is the probability of state i. Finally, the reduced uncertainty resulting from the additional knowledge of first order transition probabilities between events (postures or clusters) is given by H2=−sum[p_i [sum p_j log(p_ij)]], where p_i is the probability of i and p_ij is the probability of going to j given the current state is i.

### Mapping 10 clusters to 3 higher order clusters

Mapping the 10 clusters from the dendrogram to the squares along the diagonal in the decomposability matrix, we observe that b1, b2, b3 and b4 make up one square of postures (henceforth referred to as B), r1, r2 and r3 another (henceforth referred to as R) and g1, g2, g3 make up the third (henceforth referred to as G).

### Generating roaming

Worm roaming behavior was generated by repeatedly simulating the precise sequence of chunks or clusters: b1→ b2 → b3 → b4 → r1 → r2 → r3 → g1 → g2 → g3. Inside a particular chunk in one iteration of the above sequence, a posture in that chunk was randomly chosen and control passed onto the next chunk in the sequence, repeating the process.

### Properties of sequence of chunks

A one-to-one mapping between the 90 postures and clusters B, R and G enables us to encode worm postural sequences into sequences of B, R and G. For instance, a particular worm postural sequence could be abstracted as a sequence of frames characterized by the cluster identities, such as {B, B, G, G, G, R, R, R, B, B, R, R, R, G, G}. From such a cluster sequence, we can ascertain various properties of worm behavior based on the properties of triplets like B→R→G, or B→R→B. Cluster triplets were considered according to a sliding window that advanced one cluster at a time in the behavioral sequence. In the behavioral cluster sequence above, the triplets that formed part of the analysis were B→G→R, G→R→B, R→B→R, B→R→G. We then treat the triplets B→R→G, R→G→B and G→B→R as being equivalent and count all of them as the B→R→G triplet. Similarly, triplets of type B→G→R, G→R→B and R→B→G are counted as the same reversing B→G→R triplet.

The data used in this work (**Yemini et al., 2013; Schwarz et al., 2015**) sampled worm videos at 6 frames per second that is used to calculate the time spent in each frame (approximately 166ms) in a sequence of clusters as shown above. Time in seconds spent in each frame is added to compute the time spent in each of the identified triplets. This frame-based addition of time takes into account the repeats (same posture identified in consecutive frames) in the behavioral postural sequence. For instance, time computations on the postural sequence {1, 1, 1, 2, 3, 4, 4, 4, 5} takes into account the time spent in each of the frames even if consecutive frames have the same posture in them. Thus for that sequence, the total time taken by the worm to complete this sequence is 1.5 seconds. The assignment of sub-cluster labels to a postural sequence is done after ignoring the repeats. Again, using the sequence above as an example, repeats are first ignored such that relevant postural sequence becomes {1, 2, 3, 4, 5}, for which the sub-cluster sequence becomes {g1, g1, b1, b2, g2}. At a higher level, the sequence becomes an instantiation of the G→B→G rule. Thus, the time spent in this particular instantiation of the G→B→G rule is 1.5 seconds.

Let us now imagine a postural sequence {1, 2, 3, 4, 5, 4, 3, 2, 1} (after repeats have been removed), for which the sequences generated at higher levels of abstraction then become {g1, g1, b1, b2, g2, b2, b1, g1, g1} and {G, G, B, B, G, B, B, G, G}. According to the sliding window protocol used in all the analysis in this work, the cluster triplets whose various properties are computed are G→B→G and B→G→B. In this way, posture sequences are converted into sub-cluster and cluster sequences and the time spent in each instantiation of a behavioral rule is computed. The disruption of b1 → b2 → b3 → b4 → r1 → r2 → r3 → g1 → g2 → g3 is achieved by alternating between consecutive sub-modules. For example, a sequence such as b1 → b2 → b1 → b2 → b3 → b4 → r1 → r2 → r1 → r2 → r3 → g1 → g2 → g3 involves two alternations (b2→b1 and r2→r1) as opposed to the smooth b1→g3 sequence.

A sequence of triplets (in terms of B, R and G) when investigated at the corresponding level of 10 sub-clusters is deemed non-smooth if the number of alternations in the sequence are greater than 2. If the worm is in the regime of B→R→G rule, but satisfies either of the following conditions: (i) the underlying sub-cluster sequence is non-smooth and at least one posture is repeated more than two times, or (ii) the underlying sub-cluster sequence is non-smooth then that particular sub-sequence of postures is assigned to the “Dwell 1” rule. If on the other hand, the worm is in the regime of B→R→G rule, and satisfies the following condition: the underlying sub-cluster sequence is smooth and the number of unique sub-clusters in the sequence is greater than 5 then that particular sub-sequence is classified to belong to the b1→g3 rule (roaming rule), even though it might have less than 10 sub-clusters.

We take a heuristic of greater than 5 smooth sub-clusters in a sequence, as that means that the worm moves a relatively long distance as compared to a situation in which the sub-cluster sequence is smooth but the number of unique sub-clusters are only 2 wherein the worm actually does not move much farther from its previous position. If the worm is in the regime of rules of type B→R→B, B→G→B, R→G→R, R→B→R, G→B→G and G→R→G, then that particular behavioral sub-sequence is encoded as “Dwell 2” type rule. Finally, if the worm is in the regime of rules of type B→G→R, then a smooth B→G→R sequence involves no alternation in the sub-cluster sequence g3 → g2 → g1 → r3 → r2 → r1 → b4 → b3 → b2 → b1. In this case a non-smooth sub-cluster sequence would involve at least 2 sub-cluster transition in the opposite direction, e.g. a transition of type b2→b3 or r3→g1.

If for such a sequence of B→G→R rules, the following conditions hold: (i) the underlying sub-cluster sequence is non-smooth or, (ii) at least one posture is repeated more than once then that particular sub-sequence is classified as “Dwell 3”. Otherwise, the B→G→R sub-sequence is classified as g3→b1 or reversal rule.

To compute the proportion of all the different rules for each worm postural sequence, the number of occurrences of a particular type of grammatical rule in a particular worm behavioral sequence is computed and divided by the total number of occurrences of all the five types of grammatical rules observed in that sequence. Such a proportion for each rule was calculated for each worm and plotted as a function of the number of postures in the behavioral sequence of that worm. The number of postures are counted in the postural sequences where repeats (same posture identified in consecutive frames) have been removed. The number of postures in the postural sequence {1, 2, 2, 2, 3, 4, 4} is 4 and not 7, based on the sequence {1, 2, 3, 4} that is generated by ignoring the repeats.

Once the postural sequence of a worm’s behavior is encoded into a sequence of different grammatical rules as defined above, further properties associated with the grammatical rules like the time spent in each type of rule and the speed of the worm during each kind of rule is computed.

**Generating trajectories from postural sequences.** To compute the average speed of worm centroid, first the speed across each of the individual frames making up a postural sequence corresponding to a particular instantiation of a behavioral rule is calculated, using the model discussed in (**Keaveny & Brown, 2017**). Then the speed across all the frames is averaged to get a handle on the average speed of the worm centroid during a particular instantiation of a behavioral rule. In this way, a distribution of speeds during all instantiations of rules of a particular type is obtained. The model proposed in (**Keaveny & Brown, 2017**) is also used to obtain trajectory of worm centroid position from a sequence of postures. In the analysis comparing the time spent in roaming and dwelling type rules in mutant strains and N2 worms, for each mutant strain worm, only those N2 worms were used as controls that were imaged within 1 week (before or after) from the time the particular mutant strain worm was imaged. This is due to month to month variability in the behavior of N2 worms (**Yemini et al., 2013**).

## 3. Results

### Worm behavior represented as discrete postural sequences

As described at length in the previous section (**Materials and Methods**), we made use of the openly available *Caenorhabditis elegans* behavioral dataset that has been described previously (**Yemini et al., 2013**). Worms (N2, wild isolates, and several mutants) were picked to the centre of a patch of *Escherichia coli* OP50 on an agar plate, one at a time. The worms were allowed to habituate for 30 minutes before being tracked for a period of 15 minutes. Then, worm behavior was abstracted as a sequence of 90 posture templates (**Schwarz et al., 2015**). The behavioral sequence {5,5,5,4,4,3,3,2} is time warped as {5,4,3,2}, and the timing information of each posture is still conserved. In this way, worm foraging behavior can now be seen as a sequence of postures (**Figure 2**) with each posture having timing information, denoting the amount of time the worm stayed in that posture before moving to the next one.

**Figure 2.**
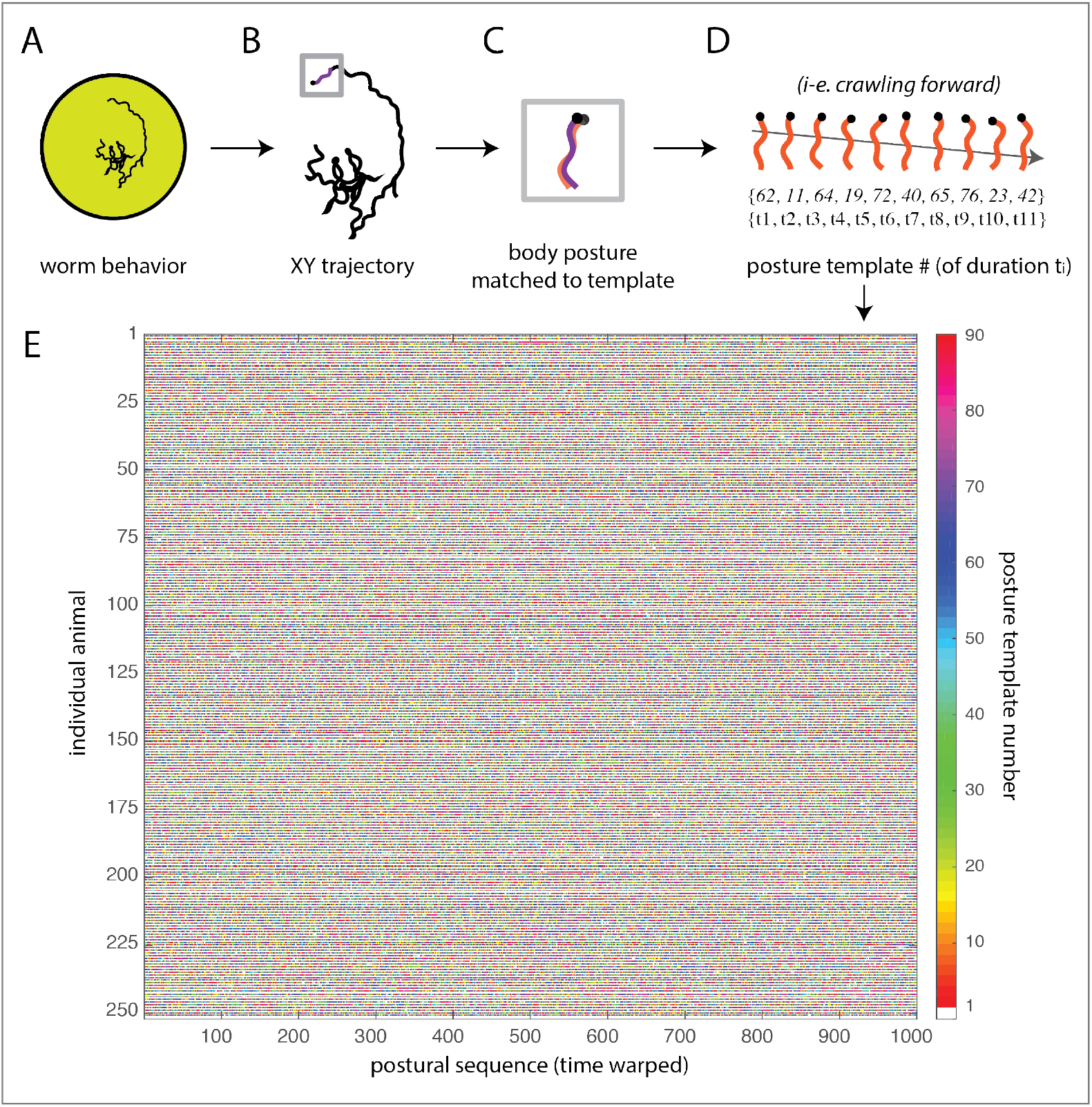
Sequencing worm behavior. (**A**) Individual worms are placed on a Petri dish to freely crawl on an agar surface with food. (**B**) Automated tracking quantitatively extracts trajectory and postural data. (**C**) Each posture shape (purple) is mapped onto one of 90 template postures (orange). (**D**) Worm behavior is then described as a sequence of discrete postures with their durations (forward locomotion example motif). (**E**) After time warping (concentrating on sequences, leaving aside durations), the behavior of hundreds of individual animals can be represented as behavioral sequences using the “alphabet” of such 90 templates (color bar).

### Postures exhibit different timescales and are flexibly used across contexts

To investigate the extent of flexibility exhibited by individual postures, based on the multiple contexts in which they occur, we plotted the probability of their recurrence after a particular time interval (**Figure 3**), for all the N2 worms pooled together. Specifically, for each posture, we calculated the amount of time elapsed between all consecutive occurrences of that posture and computed the probability of recurrence for different time intervals. We found that some postures (let us call them E-type postures) have relatively higher probability of repeating after 2 or 3 seconds (ie. 58, 10, 81) and might be related more strongly with roaming behaviors. Other postures (let us call them D-type postures) recur after very short times (i.e. 27, 20, 25), and might be involved in pause or dwelling states.

**Figure 3.**
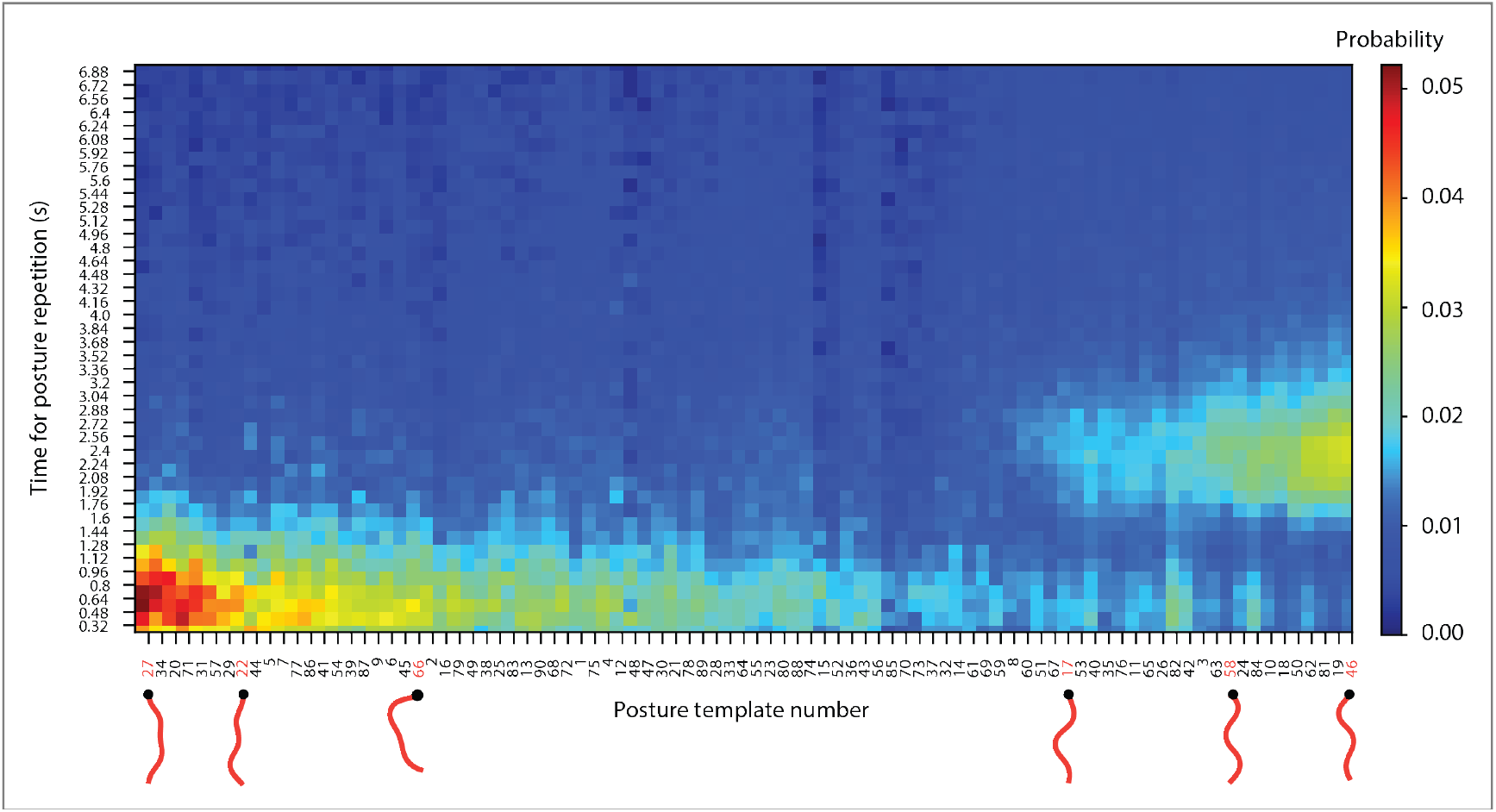
Temporal recurrence reveals two main postural uses. Probability, for each of the 90 postures to repeat after a particular time for all N2 worms pooled together. Templates are sorted such that postures having similar time profiles for repetition are grouped together, indicating candidate postures involved in dwelling (right side; x-axis) versus postures used in forward locomotion with body-wave periodicity (left side; x-axis).

In pause and dwelling states, the worm remains confined to a small region in space, and hence alternates between the same set of postures in a short span of time. In contrast, during a sustained period of roaming, a traveling wave moves along the worm’s body from its tail to the head multiple times to continuously propel it in the forward direction. The amount of time taken for the forward wave to travel along its body is usually of the order of seconds. The similarity in time of the recurrence of forward traveling wave along the worm’s body and the recurrence of E-type postures suggests that such postures may have a higher chance of being used in roaming.

Note that D-type postures also have non-negligible probabilities to recur at higher timescales (between a few seconds) and similarly E-type postures can recur at very short time scales of less than a second. This exemplifies the flexibility even at the level of primitive behavioral units like postures, where the same posture may be involved in either roaming or dwelling depending on the situation. Thus, to understand the organization of behavior in *Caenorhabditis elegans*, we need an account of this flexibility, even at the lowest level of postures.

### Worm behavior does not conform to a Markovian dynamics

We first sought to establish if worm foraging can be explained by a Markov model. A Markov model postulates that the next posture in the postural sequence is dependent on the worm’s current posture. To that end, following the work in (**Berman et al., 2016**), we looked at behavioral transition matrices at different time scales. Specifically,

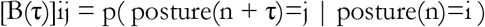

where each element of the behavioral transition matrix B, denotes the probability that the worm goes to posture j from posture i after τ discrete behavioral time steps. For example, the elements in the matrix B(1) describe the probability of moving from one posture to the next, i.e. behavioral elements that are separated by just one behavioral time step.

As shown in **Figure 4A**, when the postures are ordered in a particular way, there is a conspicuous structure in the B(1) matrix. In the forthcoming sections, we discuss the procedure used to order the matrix in this particular manner. As we increase τ from 1 to higher values, we should expect that the structure present in the B(1) matrix to progressively degrade, because as move further in time away from the current state, the ability to predict the behavioral state decreases. Alternatively, if the behavior of the worm were organized in a Markovian manner, then on should expect that:

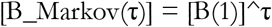

**Figure 4.**
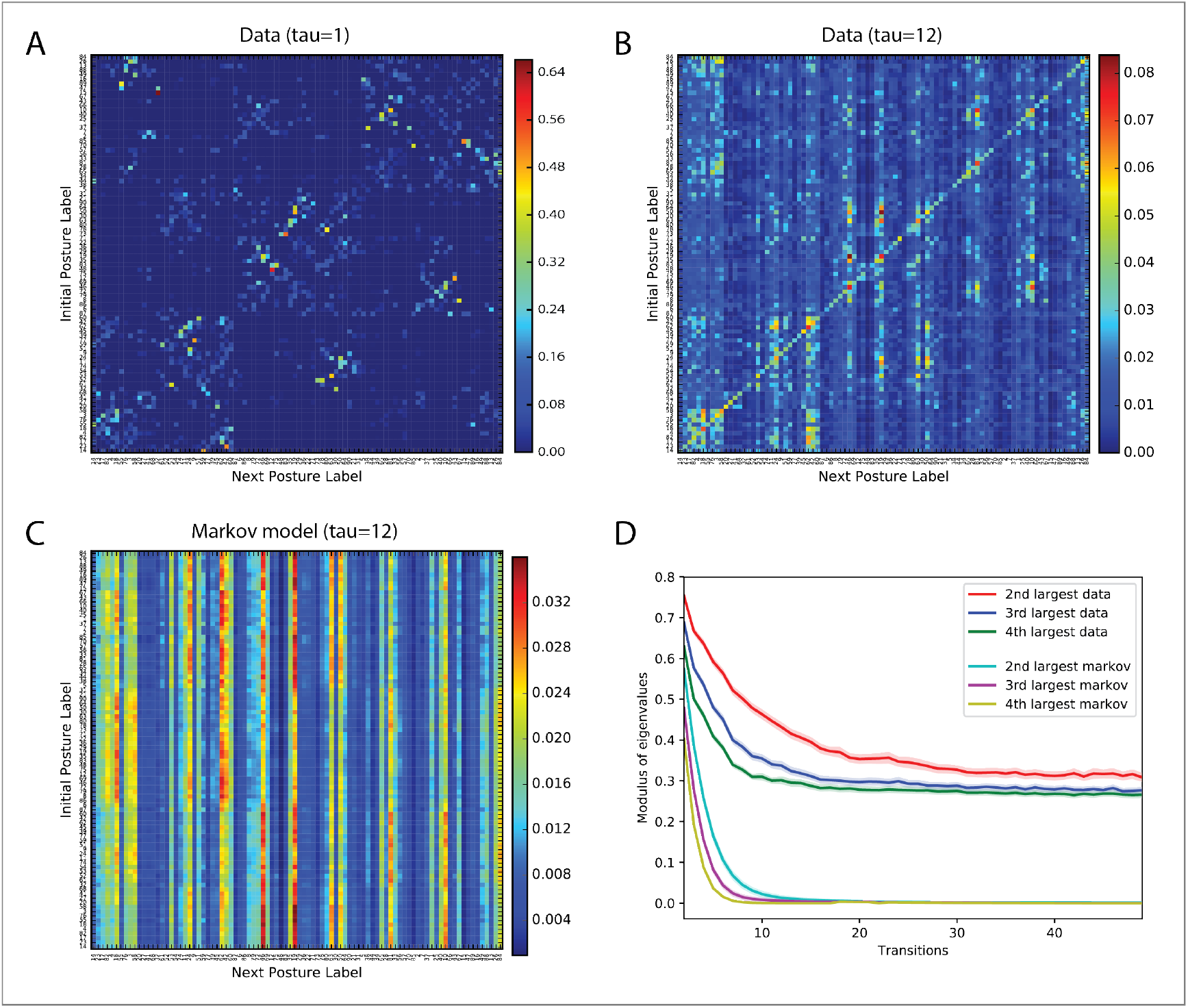
Markov transition probabilities reveal long behavioral timescales but fail to capture worm behavior. (**A**) Posture transition matrix denoting the probability of transition between pairs of postures for all the foraging N2 worms pooled together. Distinct block diagonal structure can be discerned from this matrix alone (see Figures 7 and 9, later). (**B**) Posture transition matrix looking 12 steps in the future still retains some structure, indicating that the current posture can be predictive of a posture well into the future. (**C**) Essentially all the structure is lost for the first order Markovian process if we look 12 states into the future. (**D**) Long timescales involved in worm behavior quantified by the rate of decay of the largest k eigenvalues of posture transition matrices characterized by both the Markovian process as well as actual worm behavior. The relatively slow decay in the case of actual worm behavior quantifies the intuition that actual worm behavior is modulated at higher time scales than that given by a Markovian process (solid curves denote average across all N=1287 individual worms; shaded region denotes s.e.m.).

The eigenvalues of B(1) (denoted by λi) have the property that λ1 ≥ λ2 ≥ λ3 ≥ … ≥ λn with the largest eigenvalue λ1 equal to 1. The slowest time scale in a Markovian system is governed by |λ2|, resulting in a time decay of t2 equal to −1/log|λ2|. Calculating t2 for B_Markov(1) for all the worms pooled together gives t2≈6.4 transitions. Hence, any memory that extends beyond 7 transitions would provide evidence for states that modulate behavior at a longer time scale. Visualizing B(τ) and B_Markov(τ) for τ=12 (≈2*t2) in **Figure 4B** and **Figure 4C** respectively, shows that there is some block diagonal structure that still persists in the actual behavioral data at a longer time scale as compared to a Markovian system which loses all initial structure. This intuition is quantified in **Figure 4D**, where the largest eigenvalues (leaving the largest whose value is 1) of the Markovian system and the actual data are plotted as a function of future time in terms of postural transitions. We can see that the rate at which the eigenvalues of the first order Markovian system decay as a function of time is much greater than the actual data. These analyses demonstrate that worm foraging behavior has a longer time scale than would be predicted by a first order Markov model. Note that the longer time-scales in behavior can in principle be captured by a higher order Markov model, but that model would still be limited conceptually because it assumes a single time-scale governing the overall behavior, whereas behavior is known to be modulated at multiple timescales (**Berman, 2018**).

### Substitution captures the degeneracy and reusability of behavioral sequences

Flexibility in behavior via degeneracy and re-usability coupled with hierarchical organization is nicely exemplified in the case of verbal behavior. Hierarchically organized grammatical rules between categories of words (like nouns, verbs, etc) or phrases specify the constraints according to which different sentences can be generated. Let us imagine the following grammatical rules that specify the hierarchy (see also **Figure 5**):

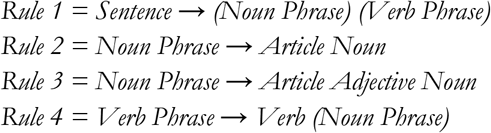

**Figure 5.**
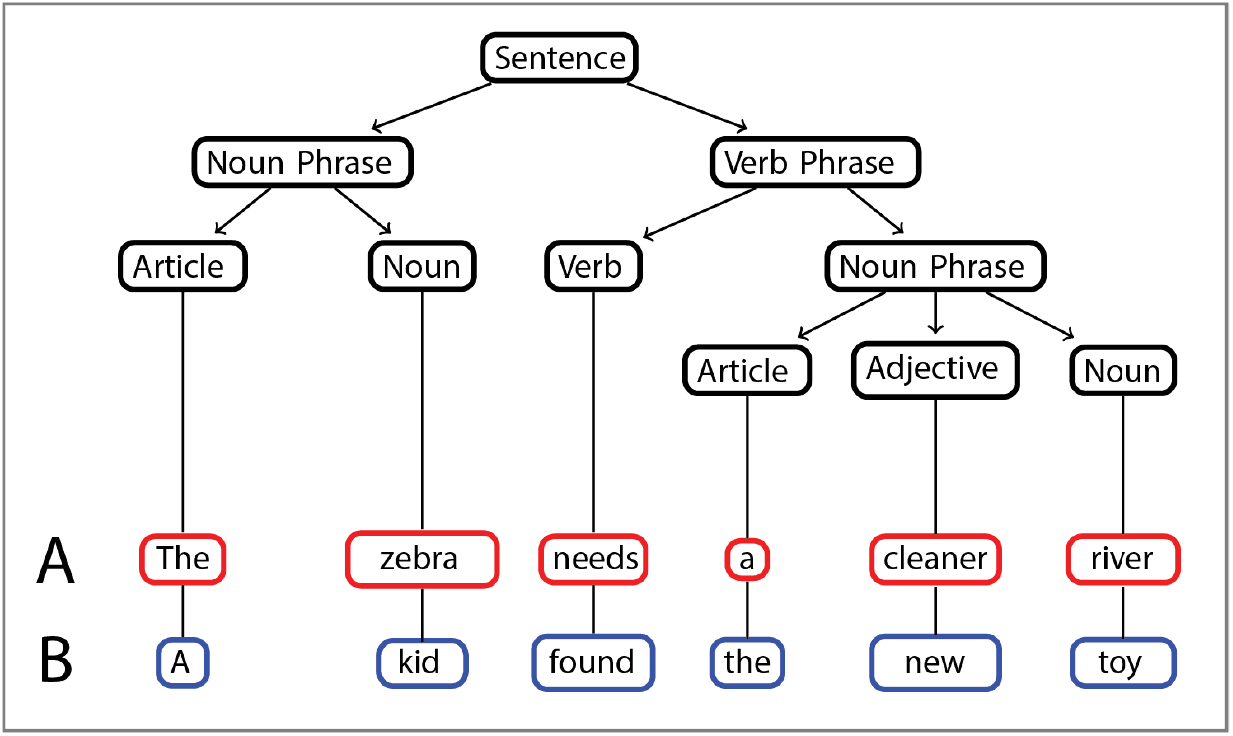
How substitution allows for flexibility within a hierarchical structure. At the lowest level, the word “zebra” can be substituted by the word “kid” to generate a new sentence (type-2 substitution), while the grammatical rules remain the same. Above the lowest level, the same description (i.e. “Noun”) can give rise to two different higher order chunks (i.e. “Noun Phrase” and “Verb Phrase”) under differing context (type-1 substitution).

It can be seen that the same lower level chunk “Noun” is re-used to generate substitutable higher level chunks in the hierarchy, “Noun Phrase” and “Verb Phrase”. We define this type of substitution, that involves re-using the same lower level description in different context and results in the substitution between higher-level chunks as Type 1 substitution. Furthermore, we also see that the same grammar can generate two different sentences by substituting words belonging to the same category like an “Article” or “Noun” (“zebra” being substituted by “kid”). We define this form of substitution that achieves degeneracy (different words belonging to the same higher order category) in a hierarchical system as Type 2 substitution. In this way, substitution can be conceptualized to capture degeneracy and reusability that lend flexibility to animal behavior.

### Behavioral modules reflect flexibility and combinatorial choice in sequence generation

Given the failure of a Markov model to account for worm foraging behavior, we explored the possibility that foraging behavior might be organized hierarchically (**Lashley, 1951; Schank & Abelson, 1977; Miller et al., 1960**). It might thus be possible to divide the worm behavioral repertoire into meaningful modules and uncover the rules of interaction between such modules (**Simon, 1962**). We hypothesized that substitution dynamics coupled with hierarchical organization, implementing degeneracy and re-usability of behavioral elements at various levels in the hierarchy might help explain the flexibility that is synonymous with behavior. Taking inspiration from the twin principles of hierarchical organization and substitution dynamics in the domain of verbal behavior, we sought modules of worm postures such that they can be used in a manner that generates flexible/variable behavioral sequences. To obtain groupings of postures from behavioral sequence data that respect variability, postures are put together in a module, if they are mutually substitutable (**Figure 6A**), with respect to their transitions to other postures. Specifically, two postures are put in the same module, if the correlation between the incoming transitions to the respective postures as well as the correlation between the outgoing transitions from them is high (see **Materials and Methods**). The procedure is illustrated with the help of an example of a restaurant menu as shown in **Figure 6B**(**Kalmus, 1969**). Given a restaurant menu where a sequence of dishes can be taken by people respecting the constraint that only one dish can be chosen from each course, eight sample dish sequences can be generated from this menu.

**Figure 6.**
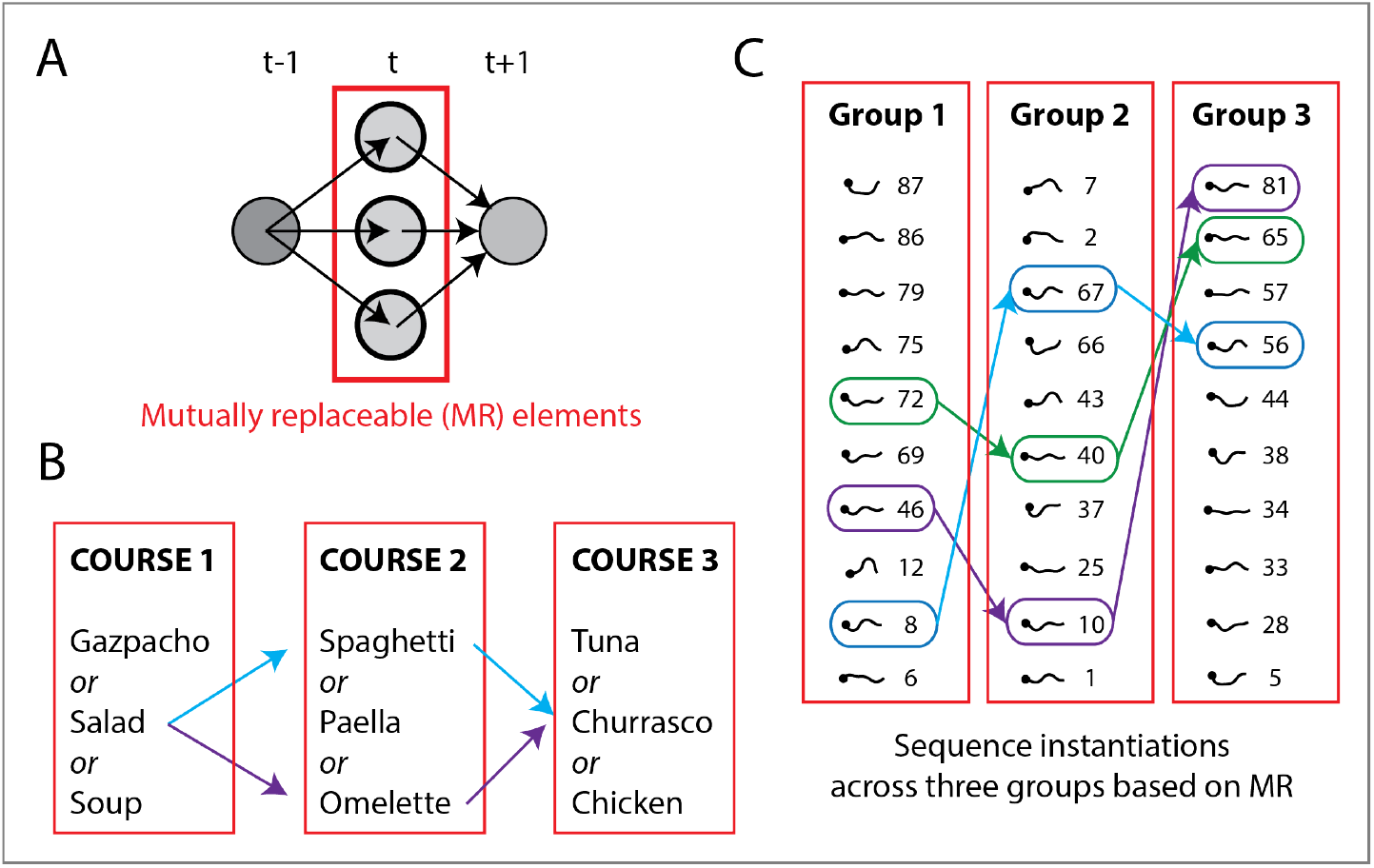
Mutual replaceability. (**A**) The essence of the algorithm: sequence elements that come from and lead to the same states are grouped together. Two examples illustrating modules comprising substitutable items and how they can generate flexible sequences: (**B**) A meal comprising a succession of three courses, each having substitutable food items; a person can choose only one item from a course leading, and so a variety of different sequences are possible. (**C**) The meal analogy made concrete for worm behavior: postural modules are formed based on MR (see Figure 7 and 8), and real worm “microscopic” sequences (using an “alphabet” of 90 postural templates) are mapped onto such “mesoscopic” modules (3 for the present illustration, but 10 in total as shown in the following Figures).

**Figure 7.**
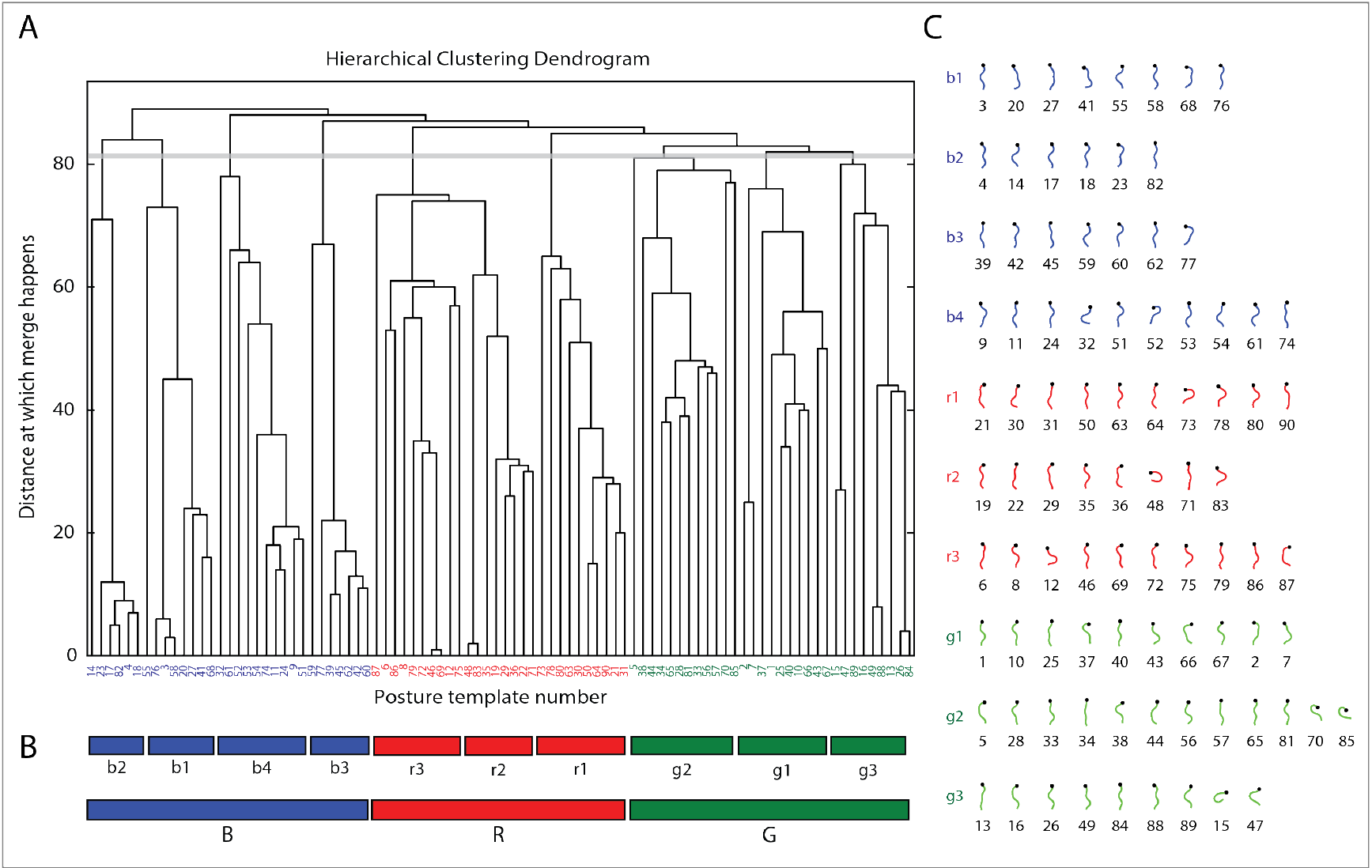
Mutual replaceability reveals 3 “macroscopic” behavioral modules made of 10 “mesoscopic” sub-modules from the 90 “miscroscopic” postural template description of worm behavior. (**A**) Hierarchical dendrogram depicting the modules and sub-modules of worm postures obtained by applying mutual replaceability. The tree is cut at a height of approximately 80 resulting in three big modules B, R and G. (**B**) Module B can further be divided into b1, b2, b3 and b4 sub-modules; R into r1, r2, r3; and G into g1, g2 and g3. Each module and sub-module consists of postures that are spanned by the spatial extent of the colored boxes. (**C**) The postural “alphabet” of each of the templates (showing explicitly the skeletons of the 90 template postures), arranged according to their belonging to each sub-module. Back to the language analogy, letters make syllables, which in turn make words and sentences.

**Figure 8.**
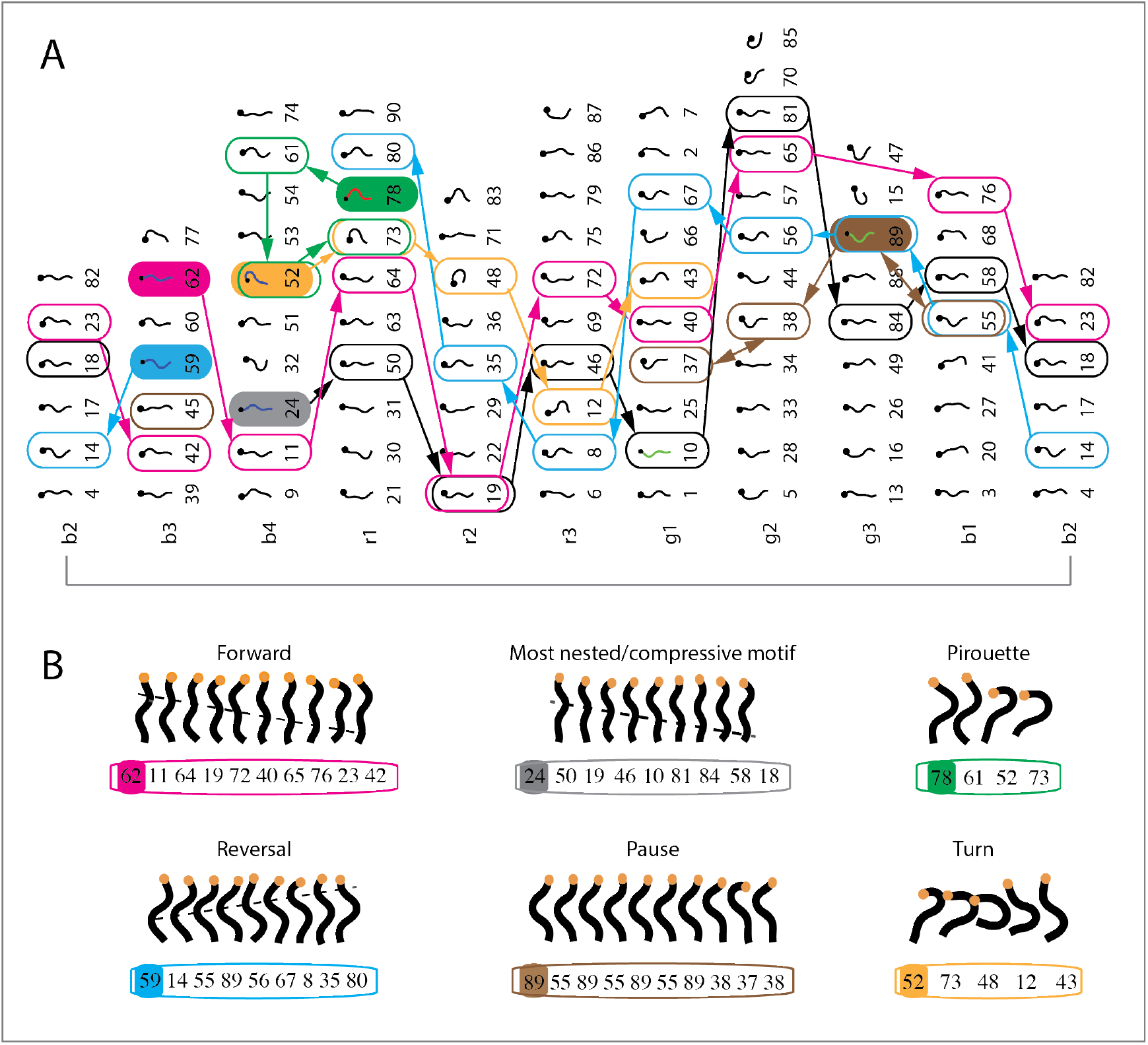
Mapping different biologically interpretable and meaningful worm postural dynamics to their behavioral modules, while establishing their rules of interaction. (**A**) The 10 sub-modules with their corresponding postural templates are arranged as subsequent columns. Posture-to-posture transitions of real data sequences are then highlighted for six typical worm behaviors depicted in (**B**), corresponding to motif sequences found in (**Gomez-Marin et al., 2016**).

**Figure 9.**
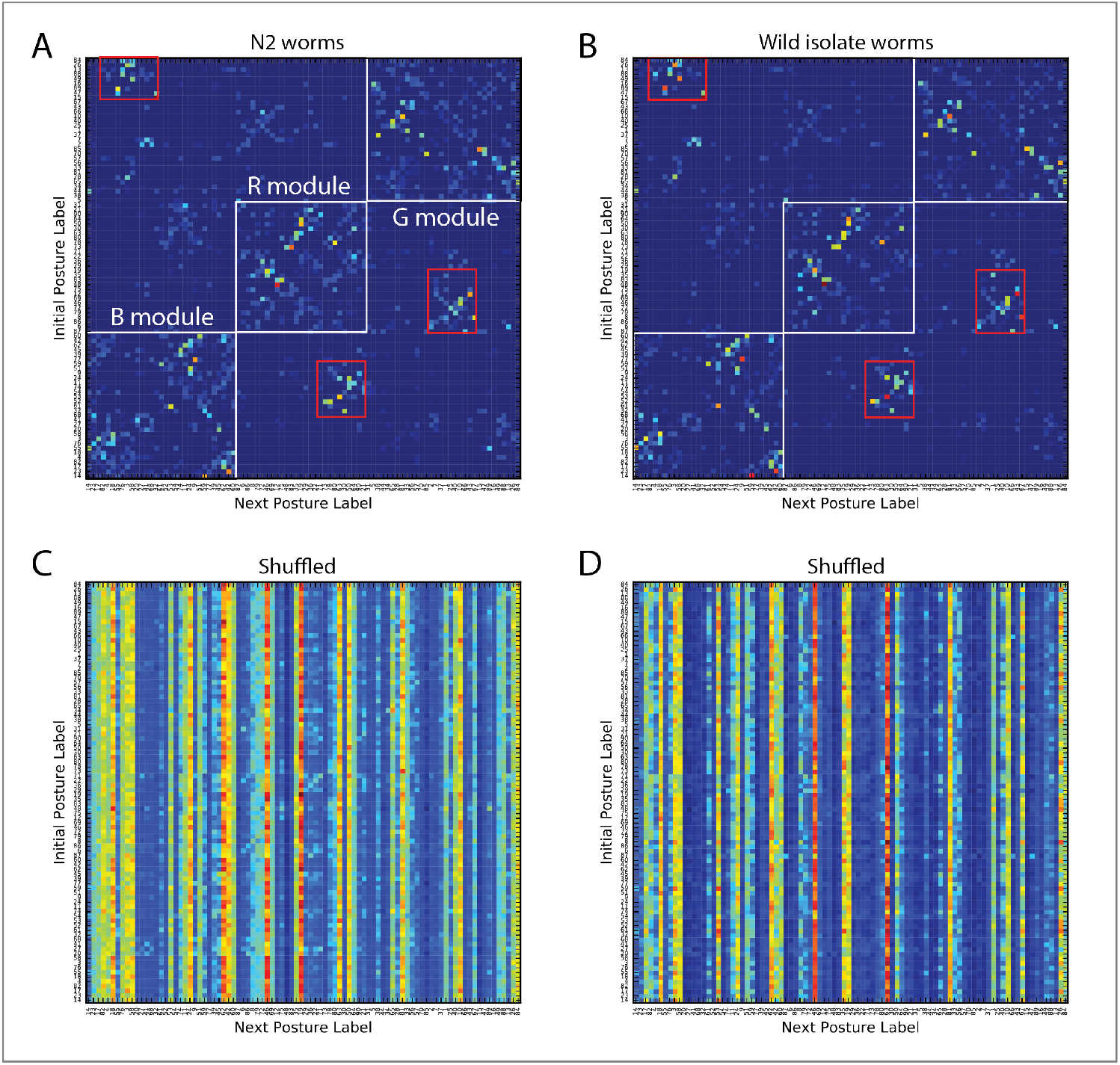
Structure emerges in postural transition matrices upon sorting based on mutual replaceability. Rearranging posture transition matrices according to the modular structure given by mutual replaceability reveals three big modules along the diagonal (white squares) with clear-cut transition structure amongst them. (**A**) Transition matrix between the 90 postures for all individuals pooled together for N2 worms, and (**B**) for wild isolate worms. The 90 postural templates are ordered according to the structure discovered by mutual replaceability, thus sorted (x & y axes) as follows: [14, 23, 17, 82, 4, 18, 55, 76, 3, 58, 20, 27, 41, 68, 32, 61, 52, 53, 54, 74, 11, 24, 9, 51, 59, 77, 39, 45, 62, 42, 60, 87, 6, 86, 8, 79, 72, 46, 69, 12, 75, 48, 83, 35, 19, 29, 36, 22, 71, 73, 78, 80, 63, 30, 50, 64, 90, 21, 31, 5, 38, 44, 34, 65, 28, 81, 33, 56, 57, 70, 85, 2, 7, 37, 1, 25, 40, 10, 66, 43, 67, 15, 47, 89, 16, 49, 88, 13, 26, 84]. Transition matrices for N2 worms (**C**) and for wild isolates (**D**) lose structure when postural sequences are shuffled.

Assuming that we do not have the course content information a priori, and are only given the observed dish sequences opted by customers, then substitutability provides a way to capture the information regarding the contents of different courses. From the sequences, we observe that “Paella” and “Spaghetti” have similar items before and after them in a meal. Thus, they can be substituted for each other and hence must be part of the same module. It must also be noted, making modules in this way based on substitution implies that elements in the same module need not occur close together in time in the observed sequence. Substitution, by capturing the combinatorial aspect of how elements are combined, leads to modules that can help generate flexible sequences like the above example. We will now make the restaurant analogy concrete in the domain of worm postures (**Figure 6C**).

We applied the method of mutual replaceability (MR) (**Maurus & Pruscha, 1973; Dawkins, 1976**) designed to capture the substitution principle, on the behavioral sequences of 1287 N2 individual worms on food. The modules and sub-modules thus formed shown in **Figure 7**, with all of the 90 corresponding postural templates plotted.

Having such modules and sub-modules at hand, as shown in **Figure 7**, we can now go back to concrete instantiations the dynamics of postural sequences and see what the modules entail, possibly starting to figure out some notion of the interactions amongst them. In **Figure 8** we make such investigation tangible by means of visual inspection of the transitions between sub-modules (**Figure 8A**) of several typical worm behaviors (**Figure 8B**). Pink and grey behavioral “motifs” correspond to instances of forward locomotion. Note how the coarse-grained sub-module description betrays a predictive advance from right to left at the level of sub-modules. The initial posture of the motif highlighted in solid color. In blue, a locomotor reversal is shown, which actually corresponds to the sub-module sequence in reverse. In brown, an example of dwelling behavior is depicted, with a dynamics involving a back-and-forth between postural templates. In yellow, the posture sequence corresponding to a turn is shown, still moving predictably in the sub-module description from right to left. Finally, a pirouette sequence is displayed in green. In sum, recasting postural sequences as a dynamics of sub-modules based on mutual replaceability betrays order without tarnishing variability.

### Mutual replaceability reveals 3 clusters & 10 subclusters regulating behavioral transitions

Using the modules obtained by applying mutual substitutability on N2 worms moving on food conditions, we plotted the first order transition matrix between postures for the N2 and wild isolate worms (see **Figure 9A** and **Figure 9B**). Actually, this is how we had ordered the 90 postures in the transition matrix according to the ordering given by **Figure 7A**. **Figures 9A** and **Figure 9B** reveal structure in the way in which postures are used to create worm behavioral sequences. Three big modules corresponding to B, R and G in **Figure 7A** are observed along the diagonal (marked out by black squares) of the matrices. The chunk or module B can further be divided into b1, b2, b3 and b4 sub-chunks, R into r1, r2, r3 sub-chunks and G into g1, g2 and g3 sub-chunks (each of which consist of a set of postures as shown in **Figure 7C**). The transition matrices demonstrate that there is a strong tendency for the worm to go from B to R and then to G (namely, “B→R→G”), using the three smaller sized modules (marked out by red rectangles) that serve as doorways. This shows that worm foraging behavior can be decomposed into 3 higher order chunks which themselves can be decomposed into 10 sub-chunks, each of which is composed of worm postures. **Figures 9C** and **9D** show shuffle controls.

### Sustained application of a simple behavioral rule generates stereotypical yet flexible forward locomotion in wild isolate worms

Next, we next sought to determine if these decomposable chunks and sub-chunks of postures have any underlying meaning for the worm by virtue of how they interact with each other. In other words, we now wish to capture all possible instances, as those shown in **Figure 8**, in the analysis framework of **Figure 9** for the sub-module transitions. Concentrating on wild isolate worms (**Figure 9B**), we then investigated the meaning of the “B→R→G” sequence and its relation to the sub-modules (“b1” through “g4”) for generating realistic worm behavior.

As a first step, we plotted the transition matrix between the 10 sub-modules (from b1 through g4) as shown in **Figure 10A**. Specifically, there is a transition counted from sub-module b1 to b2 if there is a transition from a postures belonging to b1 to a posture belonging to b2. We can see that there is a strong predictability to wild type worm foraging behavior, with the worm taking the sub-module sequence “b1 → b2 → b3 → b4 → r1 → r2 → r3 → g1 → g2 → g3” for the majority of its movement time. **Figure 10B** is a shuffle control, whose lack of structure points at the non-triviality of the structure observed in **Figure 10A**.

**Figure 10.**
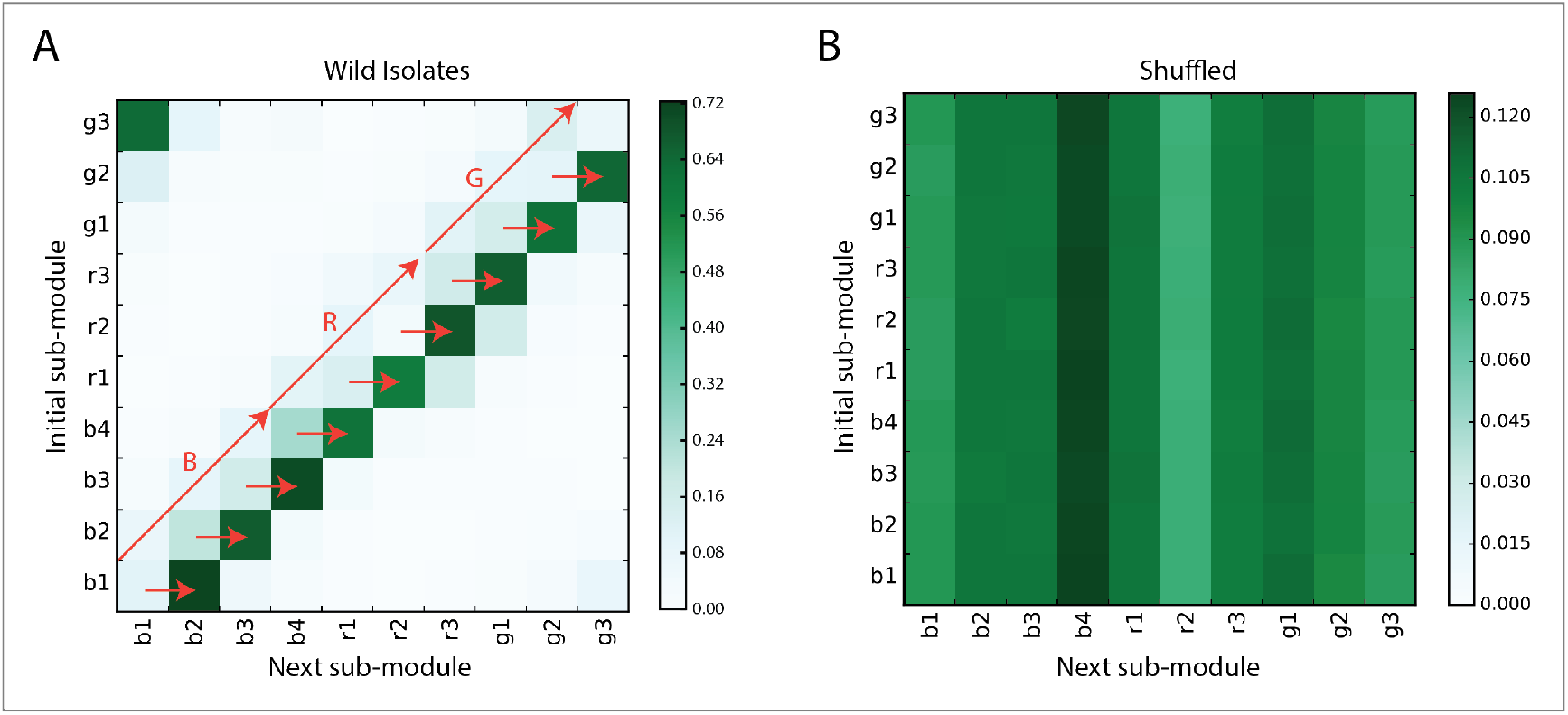
Transitions between sub-modules are stereotyped, following a cyclical rule. (**A**) Transition matrix between the 10 sub-modules for wild isolates (634 worms, pooled together). There is a very high probability that the worm moves according to the sequence “b1 → b2 → b3 → b4 → r1 → r2 → r3 → g1 → g2 → g3 → b1”. (**B**) Same as in (A) upon shuffling original sequences.

We note that our findings are qualitatively immune to changes in the number of template postures (N=90) used to capture the totality of worm locomotion. Changing the number of postures in the starting “microscopic” description of worm behavior from low (N=45) to high (N=150) shows similar patterns characterizing the transition matrix between the sub-modules (see **Materials and Methods** and **Figure S1**).

Since worms are known to perform sustained forward locomotion with limited turns and pauses for a significant amount of time, we hypothesized that the “B→R→G” rule invokes the “b1 → b2 → b3 → b4 → r1 → r2 → r3 → g1 → g2 → g3” sequence rule (or “b1→…→g3” for abbreviation) to generate roaming behavior in worms. Note that even though this is a stereotyped behavior, there is flexibility at the level of postures in the sense that whenever the worm is in a particular sub-module (say b1), it can pick any posture belonging to that particular sub-module and then move on to the next sub-module in the sequence. This flexibility results in a combinatorial explosion in the number of unique sequences that can be generated.

To test this idea of stereotypical yet flexible behavior generation, we simulated 10000 frames where each frame was represented by a worm posture. The putative forward locomotion generating sequence “b1→…→g3” was used to generate the sequence of 10000 frames. Whenever in a particular sub-module, the simulation randomly chose any one posture in that sub-module and moved on to the next sub-module to do the same. Extremely curved postures that are definitively used for making sharp turns (i.e. templates 2, 7, 70, 85, 15, 47) were not included in these simulations. Once the sequence of virtual postures was generated using the rule, we divided the sequence into 100 consecutive chunks of 100 postures (frames) each. The angles corresponding to the 48 segments of each worm posture in each of the 100 frames was then averaged across the 100 consecutive chunks to get an averaged out 100 frame chunk. The evolution of the angles corresponding to the 48 segments corresponding to each frame in the averaged out chunk of 100 frames was then visualized as shown in **Figure 11A**. The traveling wave in the forward direction (from the head to tail) shown in the figure confirms the hypothesis that sustained forward locomotion in worms is generated by the sub-module sequence “b1→…→g3” with flexibility being rendered to this stereotypic sequence by the variable choice of postures from each sub-module in every instantiation of this behavioral rule.

**Figure 11.**
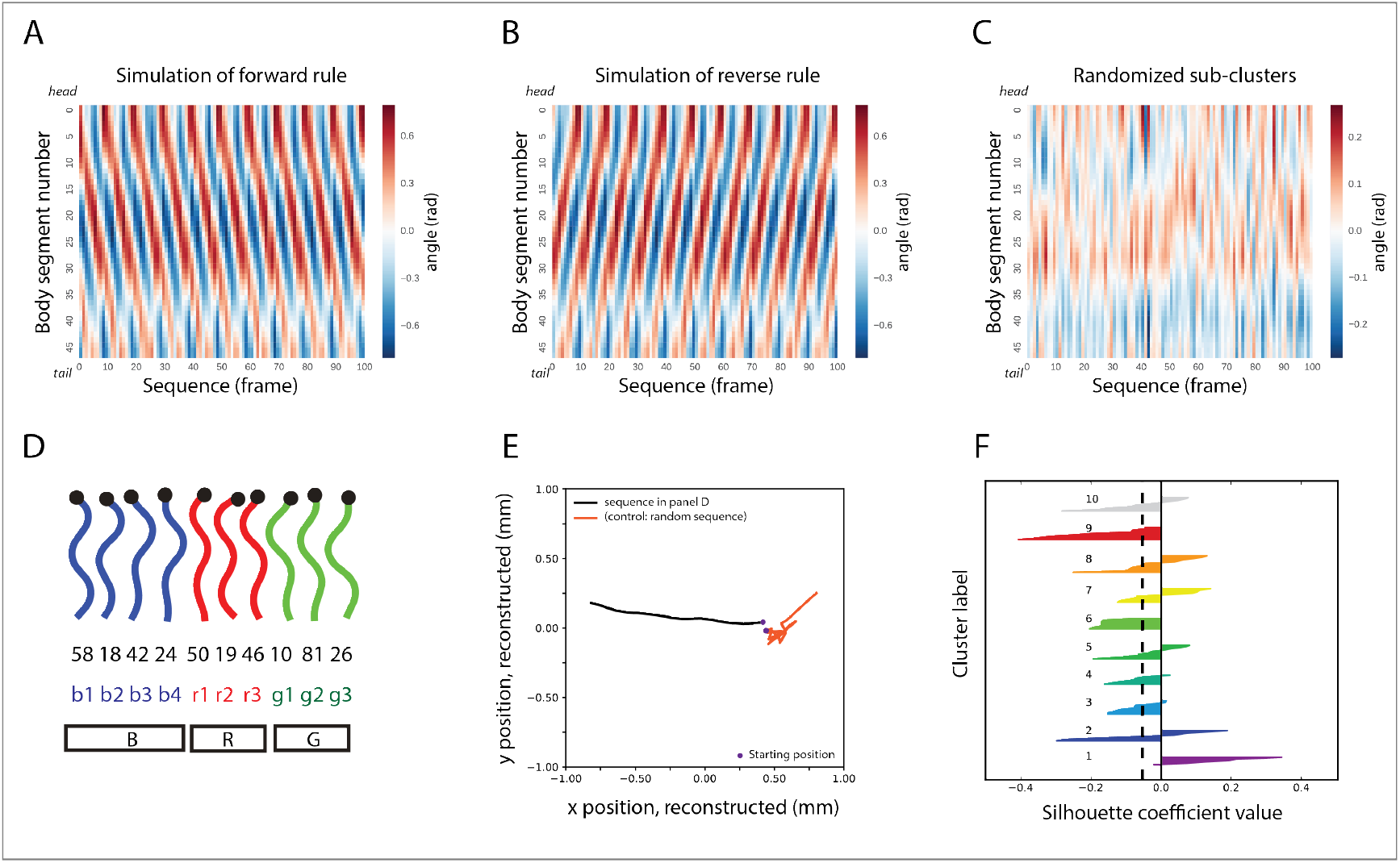
A simple rule captures and can generate roaming behavior. (**A**) Simulation of the “b1 → b2 → b3 → b4 → r1 → r2 → r3 → g1 → g2 → g3” behavioral rule (“b1→…→g3”, for short) —invoked by the hierarchically superior “B→R→G” rule— shows, on average, the progression of a traveling wave along the body of the worm that sustains forward locomotion. (**B**) Simulation of the same behavioral rule in reverse generates sustained backward locomotion. (**C**) Simulation of worm postural dynamics when the postural identities of the sub-modules remain the same but the sequence of the rule “b1→…→g3” is broken in favor of a randomized one. (**D**) Sample of a “microscopic” postural sequence from the data that respects the “b1→…→g3” mesoscopic rule. (**E**) Trajectory of center of mass position of a worm during a roaming phase characterized by 3 consecutive “b1→…→g3” sequences (as in D) comprising approximately 30 postures from an actual N2 worm (in black) and reconstructed trajectory based on application of the randomized sequence rule mentioned in (C) also consisting of 30 postures (in red). (**F**) Contribution of postural similarity to the behavioral rule. Silhouette values of all postures (ordered in a decreasing fashion) according to their membership to the 10 sub-modules generated by mutual replaceability show low intra-module morphological similarity between postures of the same cluster (black dashed line depicts average silhouette value across the 10 sub-clusters).

If “b1→…→g3” encodes smooth forward locomotion in worms, then “g3 → g2 → g1 → r3 → r2 → r1 → b4 → b3 → b2 → b1” behavioral rule should in principle generate reversal behavior. To test this idea, we simulated 100 worms each consisting of 100 frames, using the above behavioral rule, with each frame corresponding to a particular posture. Whenever in a particular sub-module, the simulation randomly chose any one posture in that sub-module and moved on to the next sub-module to do the same. Using the same procedure as was used to generate **Figure 11A**, the evolution of the angles corresponding to the 48 segments of each frame was then visualized as shown in **Figure 11B**. We can clearly see from the figure that a wave travels from the tail to the head of the worm confirming the hypothesis that the “g3→…→b1” grammatical rule indeed encodes reversal behavior.

The red rectangles in **Figure 9B** are the postures comprising sub-modules b4, r3 and g3, enabling the chain of “B→R→G” to accomplish forward locomotion. Note that multiple consecutive instances of these rules imply that, at the higher level of chunks, the “B→R→G” sequence gets instantiated multiple times like {B, R, G, B, R, G, B, R, G, B, R, G}. Due to the cyclic nature of the roaming behavioral rule, we can see that BRG roaming rule is equivalent to “R→G→ B” or “G→B→R”. During the “R→G→B” sequence, for example, instead of starting from b1, the worm would start from r1 and follow the following sequence at the level of 10 sub-chunks to complete one roaming cycle.

One important point to note is that although repeated application of “b1→g3” behavioral rule has the capacity to generate sustained forward motion, slight variations to this rule still keep the worm in roaming state. Roaming can also consist of short reversals or pauses. Roaming mostly consists of forward motion aiding the worm in traveling to farther places, but it also involves small reorientations, consisting of short reversals, as well as pauses, so that it can change direction and then travel a long distance in that direction. For example, instead of “b1 → b2 → b3 → b4 → r1 → r2 → r3 → g1 → g2 → g3”, “b1 → b2 → b3 → b4 → r1 → r2 → r3 → g1 → g2” or “b1 → b2 → b3 → b4 → r1 → r2 → r3 → g1 → g2 → g3 → g2 → g3” also encode roaming behavior. For details of heuristics used to assign sequence of postures into roaming and non-roaming states see the **Materials and Methods** section.

Finally, we plotted the trajectory of the worm centroid position from a particular sequence of postures in an actual behaving N2 worm, characterized by roaming behavioral rule using the model proposed in (**Keaveny & Brown, 2017**). As can be seen from **Figure 11E**, the trajectory of the roaming rule characterized by b1→g3, travels a far greater distance than a sequence of postures of the same length but generated by a randomized behavioral rule.

### Body morphology alone does not explain the dynamics underlying roaming behavior

One relatively straightforward way to make modules and sub-modules of postures involves clustering postures based on morphological similarity, i.e. morphologically similar postures should be grouped together into one module. To investigate the difference between sub-modules based on substitution versus those that might have been generated based on body morphology similarity, we computed the silhouette scores for all the postures based on the sub-modules given by MR (**Figure 11F**). The silhouette value for a posture p in a sub-module (given by MR) measures how morphologically similar p is to other postures in its own sub-module, as opposed to postures in other sub-modules. Large silhouette value indicates that the posture is tightly bound to other postures in its sub-module in terms of morphological similarity. The existence of high number of postures having negative silhouette value (60 out of the 90 postures) indicates that the sub-modules given by MR contain postures that are less morphologically similar to each other than compared to postures in other sub-modules (**Figure 11F**). Thus, sub-modules generated by MR capturing the substitutability between postures are different from what might be expected by modules generated based on the criterion of morphological similarity between postures.

Next, we investigated the role of body morphology in dynamics that generate worm behavior. Considering the grammatical rule for worm roaming behavior “b1→…→g3”, we looked at the morphological similarity between successive postures during all such sub-module sequences in behaving N2 worms. This morphological similarity was then contrasted with the morphological similarity that would be expected if at each posture during the “b1→…→g3” sequence, the worm transitioned to the most morphologically similar posture to the current one. We found a large difference in the distribution of morphological similarity in the roaming case versus the most similar posture next case (N=2033667, effect size, Cohen’s d = 1.102, p<0.0001, Welch’s t-test). The large effect size shows that the transitions between successive postures during roaming behavior is significantly different and cannot be captured by considering transitions between the most morphologically similar postures.

Further, it can be seen that although the profile of morphological similarity structure between successive postures is the same in “b1 → b2 → b3 → b4 → r1 → r2 → r3 → g1 → g2 → g3” as well as in “g3 → g2 → g1 → r3 → r2 → r1 → b4 → b3 → b2 → b1” behavioral rule, the behavior that these two rules generate is completely different from each other. If we assume that posture p1 is chosen from b1 and posture p2 is chosen from b2, then if we transition from b1→b2 or from b2→b1, the morphological similarity profile of the transition between p1 and p2 would remain the same. Even with the same morphological similarity structure between successive postures, the worm behavior generated by these two rules is completely different. While the former generates sustained forward locomotion (as in **Figure 11A**), the simulation of the latter behavioral rule generates sustained backward locomotion (**Figure 11B**).

These analyses reveal the primacy of the behavioral grammar in generating behavior, where identical morphological similarity transition profiles can generate different behavior based on the grammatical rule being used by the worm. These results show that even in a relatively simple organism like the nematode worm, there is structure in its behavior that goes well beyond morphological similarity between successive postures that the animal uses during behavior.

### Recapitulation: a behavioral grammar for roaming

Taken together, these results show that the “b1→…→g3” grammatical rule encodes worm roaming behavior. Thus, *Caenorhabditis elegans* roaming behavior is hierarchically organized, with type 2 substitution accounting for the flexible nature of roaming behavior (**Figure 12**). We note that worm roaming behavior is hierarchically organized not because the behavioral repertoire was divided into lesser number of chunks or modules, but because rules of interaction between such modules could be meaningfully elucidated (**Simon, 1962; Clarke & Crossland, 1985**), establishing correspondence between worm roaming behavior and compositional hierarchy. Note that at the level of higher order chunks (B, R and G), roaming behavior is only made possible by a specific rule combining these three chunks (“B→R→G”) and no other rules combining them (for example, “B→R→B”) would generate worm roaming.

**Figure 12.**
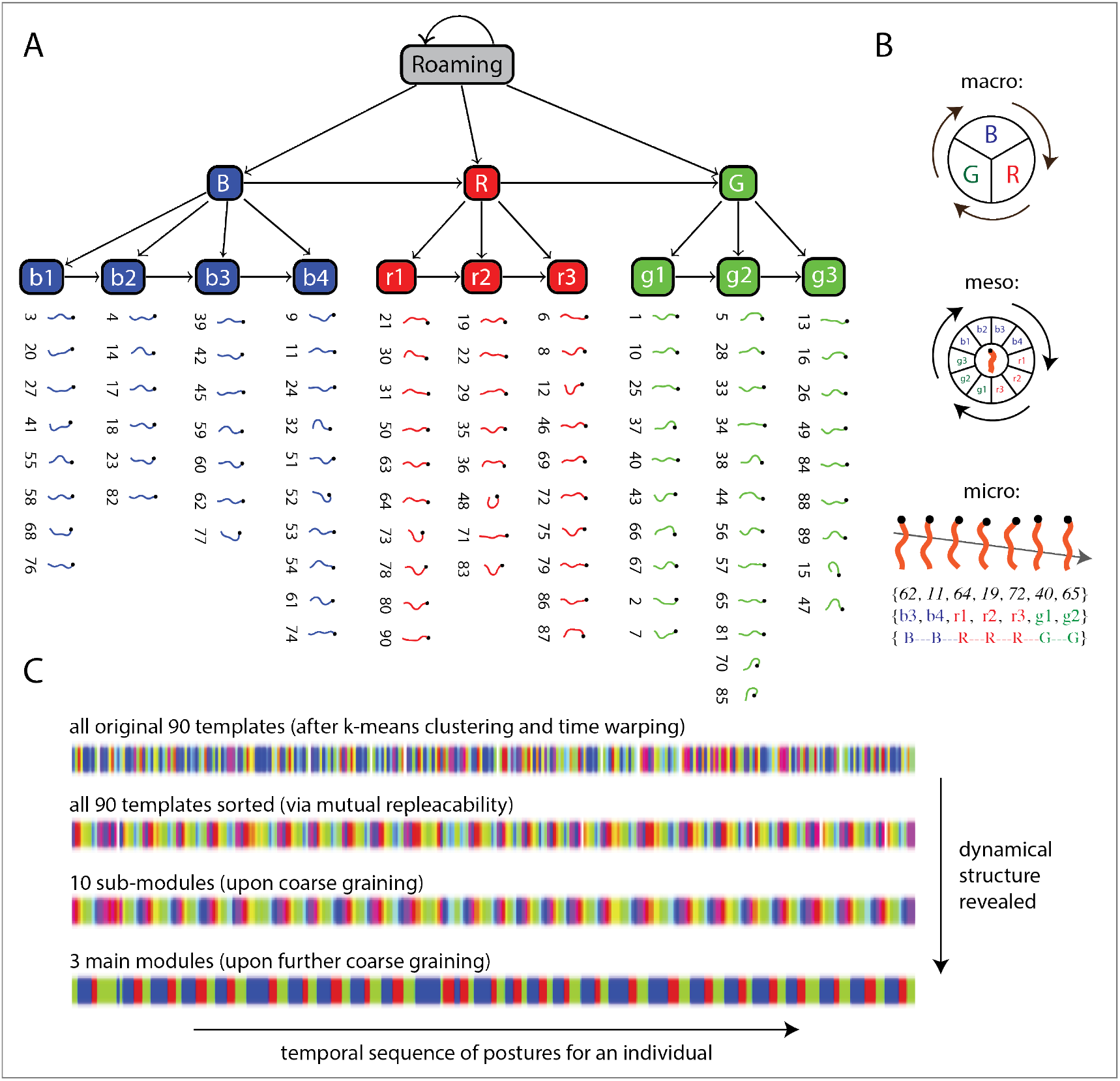
Schematic of hierarchical organization of stereotypic-but-flexible forward locomotion in worms. (**A**) Flexible order is afforded by the behavioral rules in conjunction with the possibility to choose any of the multiple postures that belong to a particular sub-module as the worm visits it (“type 2” substitution; see Figure 5). (**B**) Visual summary of the “macroscopic” and “mesoscopic” rules (module and sub-module interactions, respectively) for a “microscopic” instantiation of a postural sequence that produces forward locomotion. (**C**) How the dynamical structure of the behavior is revealed as one goes from the original 90 templates, to sorted templates, to sub-modules, all the way to modules (sample sequence data for a worm; colors depict different element types in each representation).

### Non-stereotypical dwelling disrupts the roaming rule alternating between sub-modules

In order to explain foraging behavior, it is not enough to account for the stereotypy in roaming; one must also uncover structure in dwelling despite its lack of stereotypy.

Apart from forward locomotion, *Caenorhabditis elegans* also reorients itself, by generating short reversals and interrupting forward motion frequently to change its direction. Concentrating on the behavior of N2 worms in **Figure 9A**, we first note the regions in the transition matrix that were not implicated in the generation of roaming behavior in worms, marked by yellow rectangles in **Figure 13A**, which encapsulate weaker interaction strength within themselves as opposed to the red rectangles that facilitated the “B→R→G” behavioral rule. They correspond to interactions of the type “G→R”, “R→B” and “B→G”, instead of the “B→R”, “R→G” and “G→B” transitions represented by the red rectangles.

**Figure 13.**
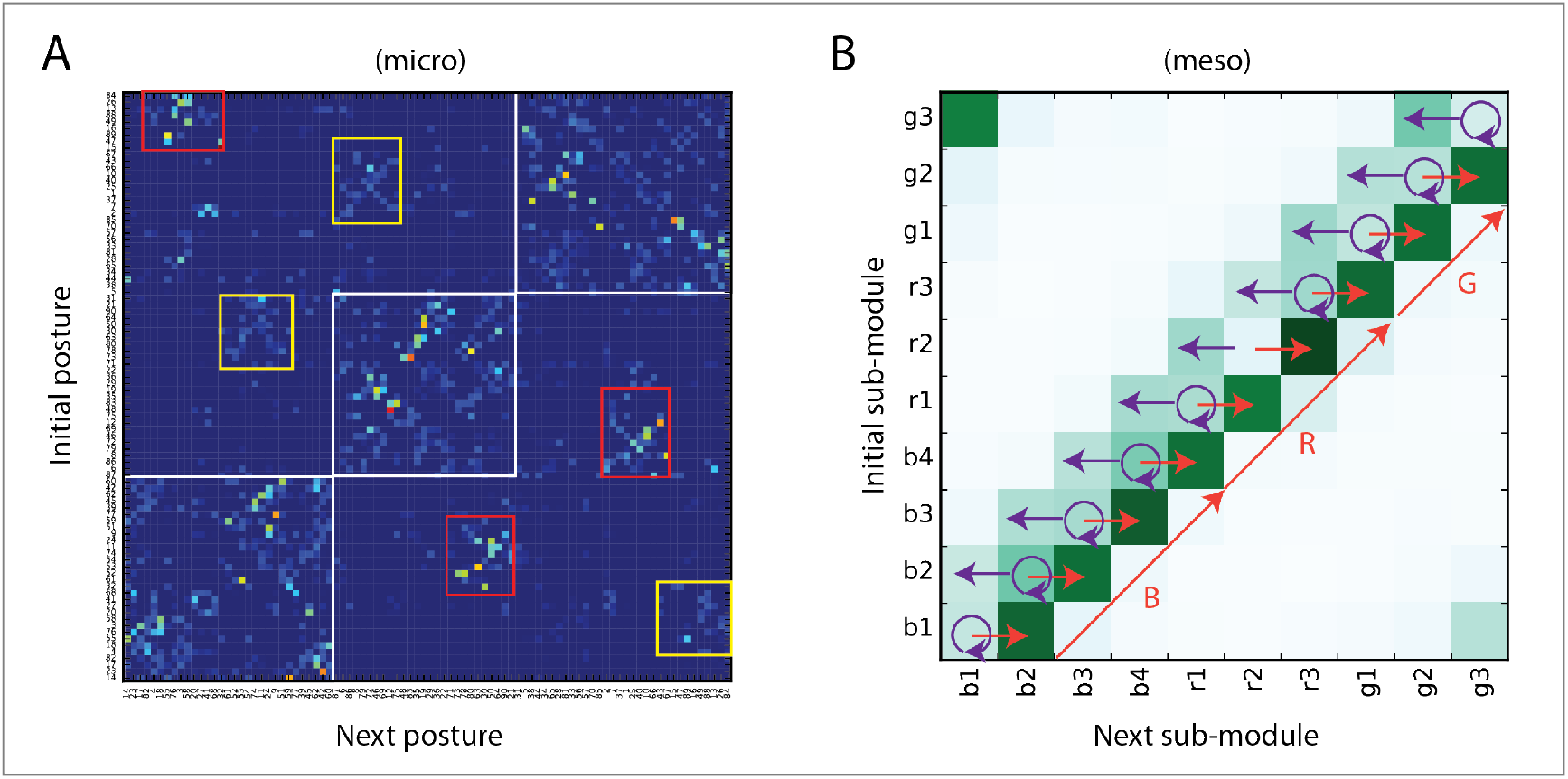
N2 worms have a higher tendency to disrupt the “b1→…→g3” rule than wild isolates. (**A**) Transition matrix for all 90 postural templates (as in Figure 9A) with the addition of yellow rectangles representing interactions between the B, R & G modules that were not implicated in forward motion. (**B**) Transition matrix between the 10 sub-modules (as in Figure 10A) for N2 worms reveals an alternation between current and previous sub-modules (purple arrows) on top of the basic “mesoscopic” sequence (orange arrows) whereby interactions between such sub-modules are increased. This means that there is an increased tendency to transition to postures of a “previous” adjacent sub-module or within the same sub-module.

Thus, comparing **Figure 13B** with **Figure 10A**, we observe that:

i. Compared to the wild type isolates, in N2 worms, there is a stronger tendency for transitions to occur between postures of the same sub-module and between postures from the current sub-module to postures of a preceding sub-module. Any sub-sequence of the “b1→g3” roaming rule must correspond to forward motion for a short time (i.e. “b3→b4→r1”). Similarly, any sub-sequence of the “g3 → g2 → g1 → r3 → r2 → r1 → b4 → b3 → b2 → b1” reversal sequence (i.e. “g1→r3→r2”) must correspond to shorter reversals.
ii. Previous work has demonstrated that worm reversals are generally associated with a decrease in their speed and form part of what is known as dwelling behavior (**Flavell et al., 2013**), that is not very stereotypic (**Gomez-Marin et al., 2016**).
iii. The full reversal sequence “g3 → g2 → g1 → r3 → r2 → r1 → b4 → b3 → b2 → b1” is not as probable as the full forward sequence “b1 →b2 → b3 → b4 → r1 → r2 → r3 → g1 → g2 → g3”.
iv. Apart from the strong tendency to follow “b1→g3” rule (similar as in wild isolates), N2 worms also display behavior that disrupts this “b1→g3” rule more strongly than the wild isolates. This disruption is very specific in the sense that it involves an increased use of more than one posture from the current sub-module that the worm is in and/or postures belonging to the preceding sub-module in the “b1→g3” sequence. It is not the case, for example, that b1 suddenly starts making increased transitions to r3 to break the “b1→g3” sequence.
v. This results in the disruption of the “b1→g3” sequence rule by alternating between consecutive submodules in the “b1 → b2 → b3 → b4 → r1 → r2 → r3 → g1 → g2 → g3” rule.

Note that disruption in the “b1→g3” sequence, in terms of higher-level modules (B, R and G) can be achieved in the following three ways, which we define as:

1. “Dwell 1”: alternating between sub-modules but still maintaining the “B→R→G” rule.
2. “Dwell 2”: achieving alternation between sub-modules by alternating at the level of higher modules by adopting rules of the form “B→(R or G)→B”, “R→(B or G)→R” and “G→(B or R)→G”.
3. “Dwell 3”: alternating between sub-modules such that the smooth reversal sequence (without alternations) of “g3 → g2 → g1 → r3 → r2 → r1 → b4 → b3 → b2 → b1” is disrupted while still maintaining the “G→R→B” sequence seen in smooth reversals.

### Timing aspects of the sequencing rules

If the behavior rules corresponding to Dwell 1, Dwell 2 and Dwell 3 encode dwelling behavior in worms, then the proportion of such rules should be higher when the sequence length of the worm during the 15 minutes recording is shorter. This is because during dwelling, the speed of the worm is slower, thus spending a greater amount of time per posture leading to a decrease in the number of unique postures that make up the worm’s behavioral sequence. During roaming, the worm can be presumed to spend relatively less amount of time in each posture due to its higher speed, thereby generating a behavioral sequence that is comparatively longer in length.

We therefore plotted the proportion of purported dwelling and roaming type behavioral rules as a function of the sequence length (**Figure 14A** and **Figure 14B**) and found that the proportion of Dwell 2 type rules decreases sharply as the sequence length increases. This decrease in the proportion is offset by a corresponding increase in the roaming type rules as the sequence length increases, lending support to the idea that Dwell 2 type rules encode dwelling behavior. In contrast, the proportion of Dwell 1 and Dwell 3 is relatively stable at a low value.

**Figure 14.**
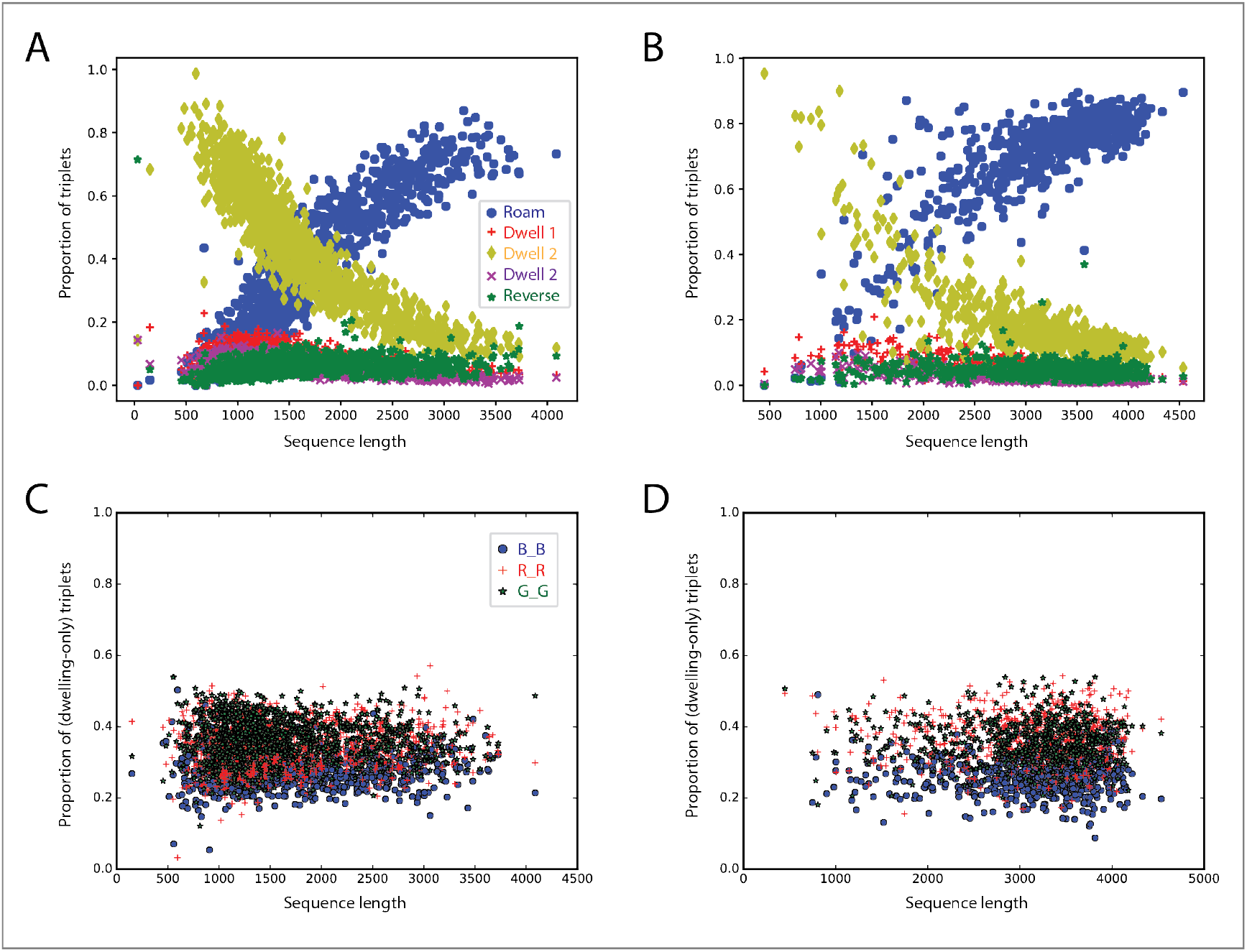
Usage of roaming and dwelling rules varies as a function of sequence length, which reflects time. (**A**)The proportion of the roaming-type behavioral rule increases as a function of postural sequence length, while the usage of “Dwell 2” rules decreases. The proportion of “Dwell 1”, “Dwell 3” and “reverse” rules remains stable. Data corresponding to all N2 worms. (**B**) The same pattern is observed for wild isolates. Note that every worms as tracked during the same amount of time (15 minutes) and that after time warping (see Figure 2) the length of their postural sequences (total number of postures, without repeats) is inversely proportional to the average time spent in each posture before transitioning to another. Thus, the shorter the sequence length, the higher the speed of alternation between different postural templates. (**C**) The proportion of the three types of “Dwell 2” type rules is similar across varying posture sequence lengths in N2 worms and also (**D**) in wild isolates (**** = p < 0.0001, Welch’s t-test; effect size: d = Cohen’s d. Violin Plots contain box plots that show the interquartile range).

As the worm moves faster during roaming than during dwelling (**Flavell et al., 2013**), we hypothesized that the speed of the worm during the “b1→…→g3” rule should be higher than during the rules hypothesized to underlie dwelling. As a corollary, we reasoned that the time spent during a single instantiation of the roaming rule ought to be lesser in comparison to the time spent during a single instantiation of dwelling type rules. Hence, we calculated the average speed and total time taken during each instantiation of all the behavioral rules (“b1→…→g3”, “g3→…→b1”, Dwell 1, Dwell 2 and Dwell 3). We averaged the speed across all the frames belonging to a particular instantiation of a behavioral rule to get a handle on the average speed of the worm centroid during a particular postural sequence given by a particular behavioral rule. Moreover, the distribution of Dwell 2 type rules shows no preference, as we found them to be nearly equally distributed (**Figures 14C** and **14D**).

It can be seen from **Figure 15A** that the average speed in N2 worms during instantiations of the hypothesized roaming rule is higher than the average speed during all instantiations of the various rules hypothesized to underlie dwelling. It is also worth noting that the speed during instantiations of the hypothesized rule for smooth reversal (“g3→…→b1”) is also quite high. The same holds for wild isolates (**Figure 15B**).

**Figure 15.**
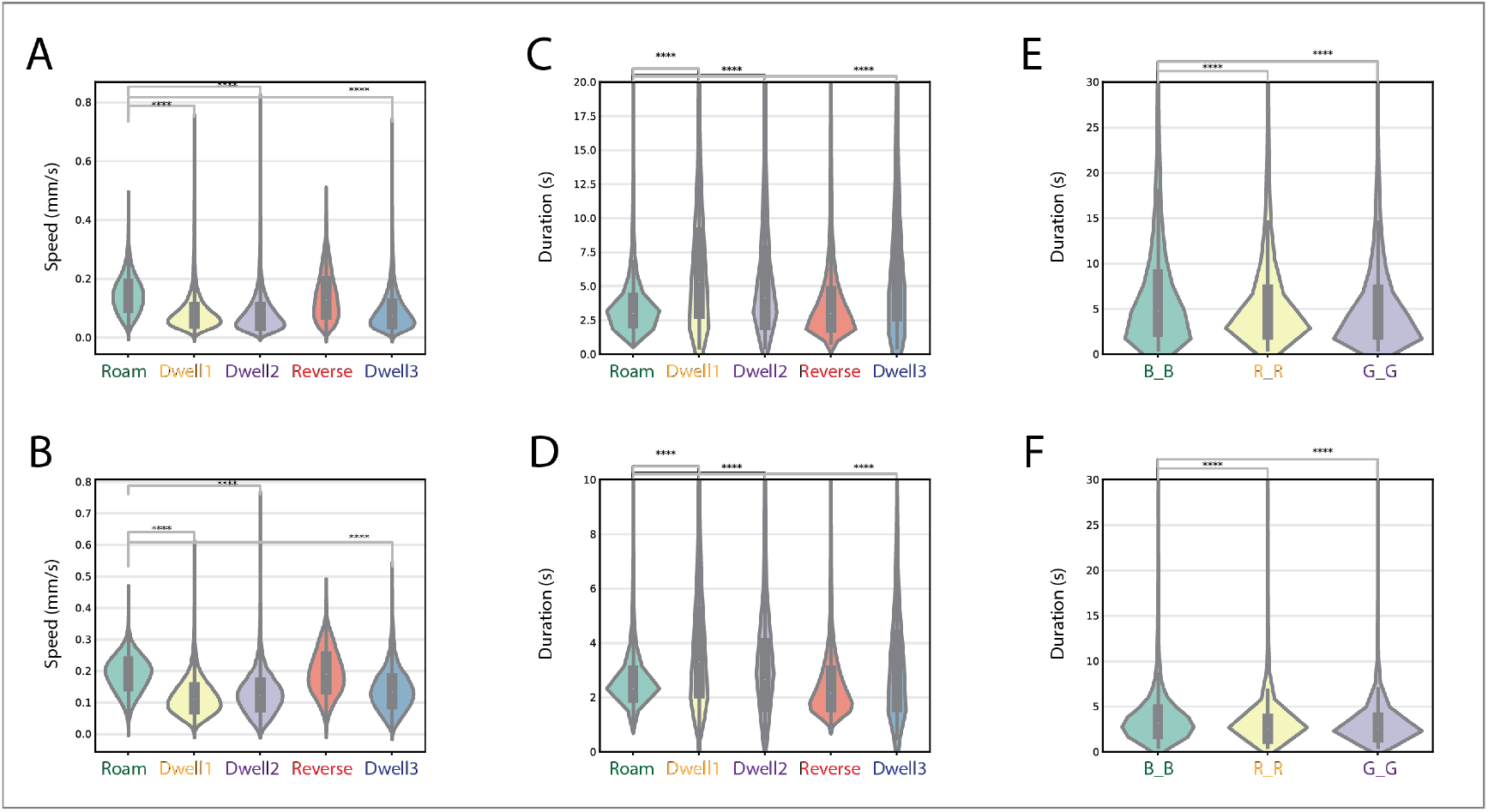
Speed and duration reflect different functional roles of roaming and dwelling grammatical rules. (**A**) The average speed of the center of mass position of the worm during instantiations of the roaming type rule is considerably higher than during the behavior rules corresponding to dwelling. Violin plots show the distribution of average speeds during each instantiation of all types of rules across all the worms (d(roam,dwell1) = 0.978, d(roam,dwell2) = 1.1, d(roam,dwell3) = 0.91; N(Roam)= 267217, N(Dwell 1)= 51538, N(Dwell 2)= 240267, N(Reverse)= 39424, N(Dwell 3)= 33287). Analyses for N2 worms and (**B**) for wild isolates (d(roam,dwell1) = 1.29, d(roam,dwell2) = 1.1, d(roam,dwell3) = 0.89; N(Roam)= 424860, N(Dwell 1)= 26492, N(Dwell 2)= 96763, N(Reverse)= 27371, N(Dwell 3)= 13105). (**C**) The total time spent during roaming-type rules is much smaller than that spent during dwelling type rules. Violin plots show the distribution of times taken to complete each instantiation for all the behavioral rules across all worms (d(roam,dwell1) = −1.05, d(roam,dwell2) = −0.51, d(roam,dwell3) = −1.19). Analyses for N2 worms and (**D**) for wild isolates (d(roam,dwell1) = −1.26, d(roam,dwell2) = −0.4, d(roam,dwell3) = −0.86). (**E**) Deconstructing the time associated with dwell 2 type behavioral rules (N(B_B)=68986, N(R_R)=83038, N(G_G)=88243; d(B_B,R_R)=0.17, d(B_B,G_G)=0.18). Again, analyses for N2 worms and (**F**) for wild isolates (N(B_B)=25588, N(R_R)=36236, N(G_G)=34939; d(B_B,R_R)=0.18, d(B_B,G_G)=0.17).

Next, as shown in **Figure 15C**, the total time spent during instantiations of roaming type rule is considerably lesser than the time spent during instantiations of rules implicated in dwelling. Specifically, the total time spent in each instantiation of all the different types of behavioral rules across all worms was computed and their distributions were then plotted. The slower speed and the higher time duration for rules of type Dwell 1 and Dwell 3 also establish their role in dwelling behavior. Again, this holds for wild isolates too (**Figures 15D**).

Since the proportion of rules of type Dwell 1 and Dwell 3 remains relatively stable across a variety of postural sequence length (**Figure 14A** and **14B**), we decided to further look into the more dynamic Dwell 2 type dwelling rules. **Figures 1E** and **15F** show that the time taken during each of the rules of type “B→(R or G)→B”, “R→(B or G)→R” and “G→(B or R)→G “is consistently higher than the time taken during the rule that characterizes roaming. Also, the timing difference between rules of type “B→(R or G)→B”, “R→(B or G)→R” and “G→(B or R)→G” amongst themselves is not significantly different (low effect size as shown in **15E** and **15F**; p-values denote significance but that is likely due to the large values of N for these comparisons).

### A brief note on predictability

To quantify the predictability afforded by dividing the 90 postures into 10 sub-modules, we calculated the H0, H1 and H2 entropies (see **Materials and Methods**) values for the two conditions (90 postures versus 10 sub-modules). H0 denotes the uncertainty in predicting the next event (posture/sub-module), if all the events are equi-probable, H1 denotes the reduced uncertainty afforded by the knowledge of individual event probabilities and H2 denotes the reduced uncertainty afforded the knowledge of first order transition probabilities between the individual postures. The values of H010, H110 and H210 (where the superscript denotes the 10 sub-modules) were 3.32, 3.31 and 2.02 bits respectively, as compared to the H090, H190 and H290 values of 6.49, 6.19 and 3.122 respectively for the 90 postures.

The reduction in entropy values for the 10 sub-module scenario indicates that we can achieve greater predictability by dividing up the 90 posture into 10 sub-modules. Note that this analysis might appear a bit non-informative because the entropy is bound to decrease once you decrease the number of objects under consideration. The counterpoint is that the way in which we decompose the 90 postures into sub-modules and then specify the dynamics characterizing interaction between them leads to functionally relevant worm behavior, thus indicating the significance of both the sub-modules and the dynamics operating on top them.

### Recapitulation: a behavioral grammar for dwelling

Taken together, these observations indicate that even in worm dwelling behavior that is thought to be relatively less stereotypic than roaming behavior (**Gomez-Marin et al., 2016**), there is predictability owing to the hierarchical nature of behavioral organization. Specifically, if we know that the worm is in dwelling state, we know that the smooth “b1→…→g3” sequence rule is broken. Furthermore, if we further know that inside the dwelling phase, the worm is in B state, then we know for sure that either “B→R→B” or “B→G→B” has to hold. Additionally, if we further know that the worm is in “B→R→B”, we can be sure that the scaffold of sub-modules that the worm will execute. In this way, the worm dwelling behavior, like its roaming behavior, is predictable yet flexible. The proposed grammatical rules are summarized in **Figure 16A** and **16B**.

**Figure 16.**
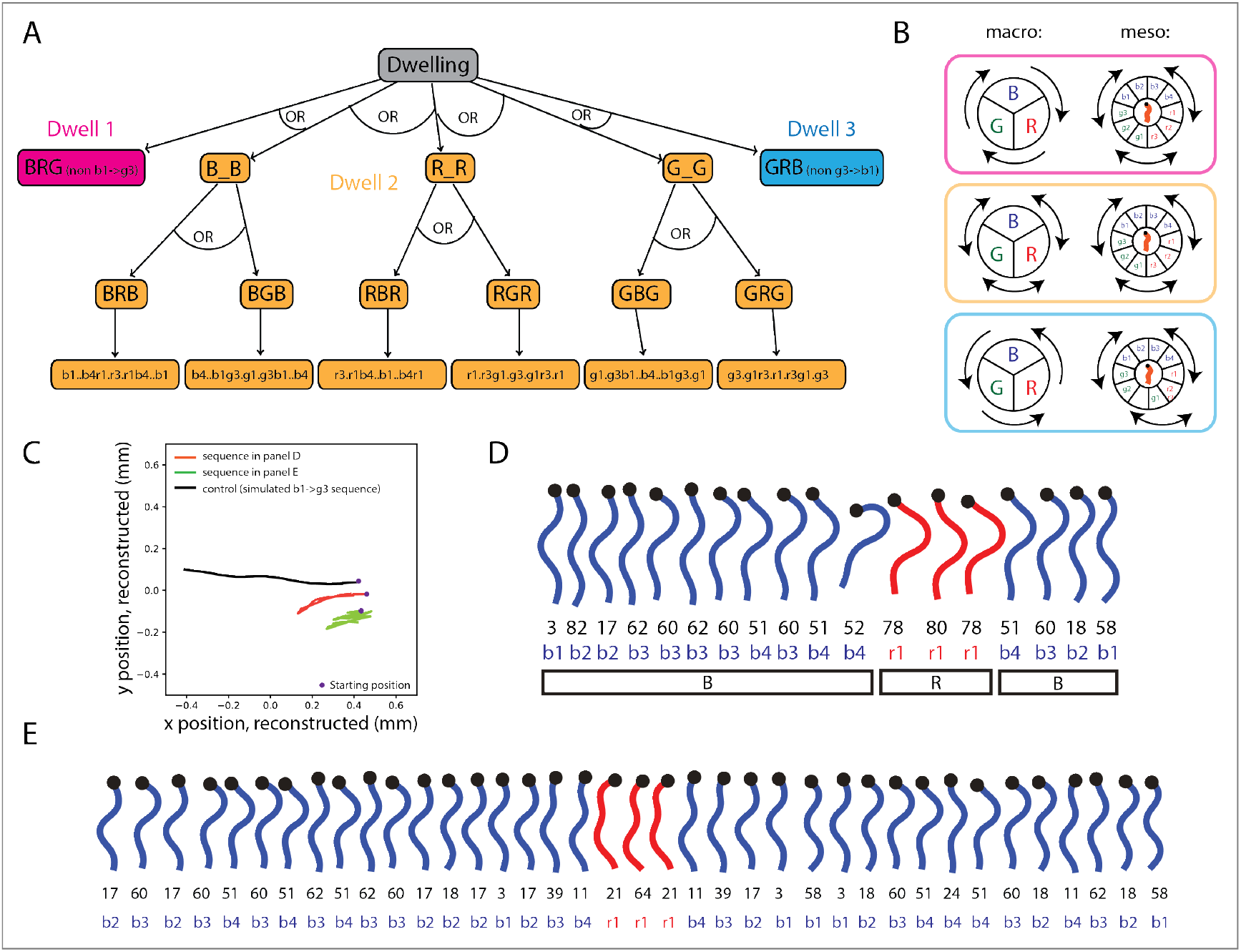
Worm dwelling behavior as a set of grammatical rules. (**A**) Schematic of dwelling behavior with the three higher Dwell 1, Dwell 2, and Dwell 3 rules. The multiple rules for realizing dwelling lend it a relatively less stereotypic character than roaming behavior. There is Type 2 substitution at the level of postures as well as at the level of sub-modules, where the different sub-modules can be involved in the generation of same dwelling behavior. (**B**) Visual summary representation of the interactions between modules and sub-modules that correspond to the dwelling rules. (**C**) Reconstructed trajectory of the worm centroid for the data posture sequences shown in (D) and (E), in red and in green respectively. As a control, we plot the trajectory (in black) produced by a roaming sequence (2 consecutive “b1→…→g3” rules) involving roughly the same of postures as in the other sequences. (**D**) A sample sequence (rule type BRB) taken from an actual foraging N2 worm. The posture sequence produces brief and minor forward locomotion frequently interspersed with backward locomotion, characteristic of dwelling behavior. (**E**) Another sample of BRB sequence with many posture alternations back and forth, showing that the behavior can be arbitrarily long while still conforming to the grammatical constraints for dwelling.

The sub-module sequence (like the lowest part of the tree in **Figure 16A** generated by the “B→R→B” sequence) provides a scaffold with sub-sequences that could get arbitrarily long while still maintaining the structure imposed by the rules. This capacity of memory is a property of hierarchical systems where the time spent in a sub-chunk inside a bigger chunk can extend to arbitrary time scales as highlighted by the posture sequence in **Figure 16E**. It shows a dwelling sequence of type “B→R→B”, like in **Figure 16D**, with almost the same set of postures making up the two sequences (see **Figure 16C**). Yet, the posture sequence in **Figure 16E** is considerably longer than the other one. Hierarchical organization that treats “B→R→B” as one single unit permits sequences where the posture sequence generated by the first B chunk inside the “B→R→B” unit can be arbitrarily long, still having memory to generate posture sequences from the R and B chunks after finishing the posture sequence from the first B chunk to make up the “B→R→B” unit. On the other hand, linear models can only look so far back in time and usually find it difficult to handle long-term dependencies in sequential data.

Thus, worm roaming and dwelling behavior can be described in terms of interaction rules between the 10 sub-modules, the interactions between which are in turn dictated by the interaction rules between three higher order chunks B, R and G, leading to predictability and flexibility simultaneously and pointing to a hierarchical organization of behavior. Note that type 1 substitution comes into the picture when we view the whole worm foraging behavior from a level of description that comprise both roaming and dwelling behavior. At that level of description, there is type 1 substitution because the same sub-module, let’s say b1 can be involved in roaming or dwelling (using roaming or dwelling specific rules) based on the context. At a higher level of abstraction, the chunk B can either be used during roaming with the invocation of the “B→R→G” sequence rule while the same B chunk can be used during dwelling through the “B→G→B” and other sequence rules.

This is the essence of hierarchical organization where the transition probabilities between chunks does not remain constant, instead depends on the higher order unit which subsumes the chunks and sub-chunks. Similar argument holds for the 10 sub-chunks b1 to g3 and finally for the 90 postures, with the same posture being used during roaming and dwelling phase with different probabilities. These roaming and dwelling specific interaction rules between chunks and sub-chunks give rise to a grammar of worm foraging.

### Proposed grammatical rules capture worm behavior in off food environments

When transferred from an environment with food to an environment without food, the worm initiates area restricted search for the first few minutes, initiating a lot of turns and staying in a small area expecting to find food nearby (**Hills et al., 2004; Calhoun et al., 2014**). But after a period of about 15 minutes, the worm switches to a dispersal behavior, whereby its rate of turns is reduced and instead it travels in extended trajectories exploring vast areas. Thus, we next asked if our proposed worm foraging grammar can capture these subtleties of worm behavior. Specifically, if our proposed worm grammar is correct, then the amount of time spent in the grammatical rules corresponding to roaming should be considerably higher than the amount of time spent in dwelling type grammatical rules, in off food environment. This is because the data that we use includes a waiting time of 30 minutes after transferring worms from an on food environment to one without food before tracking their behavior. And hence, we should expect that the worm has already finished its area restricted search (akin to dwelling) and has initiated its dispersal behavior (akin to roaming).

As shown in **Figure 17A**, the amount of time spent by N2 worms in roaming type rules is considerably higher than the time spent in dwelling type rules, in off food conditions. Also shown in the same figure is the complete opposite behavior shown by N2 worms on food where the amount of time spent dwelling is considerably higher than the time spent roaming.

**Figure 17.**
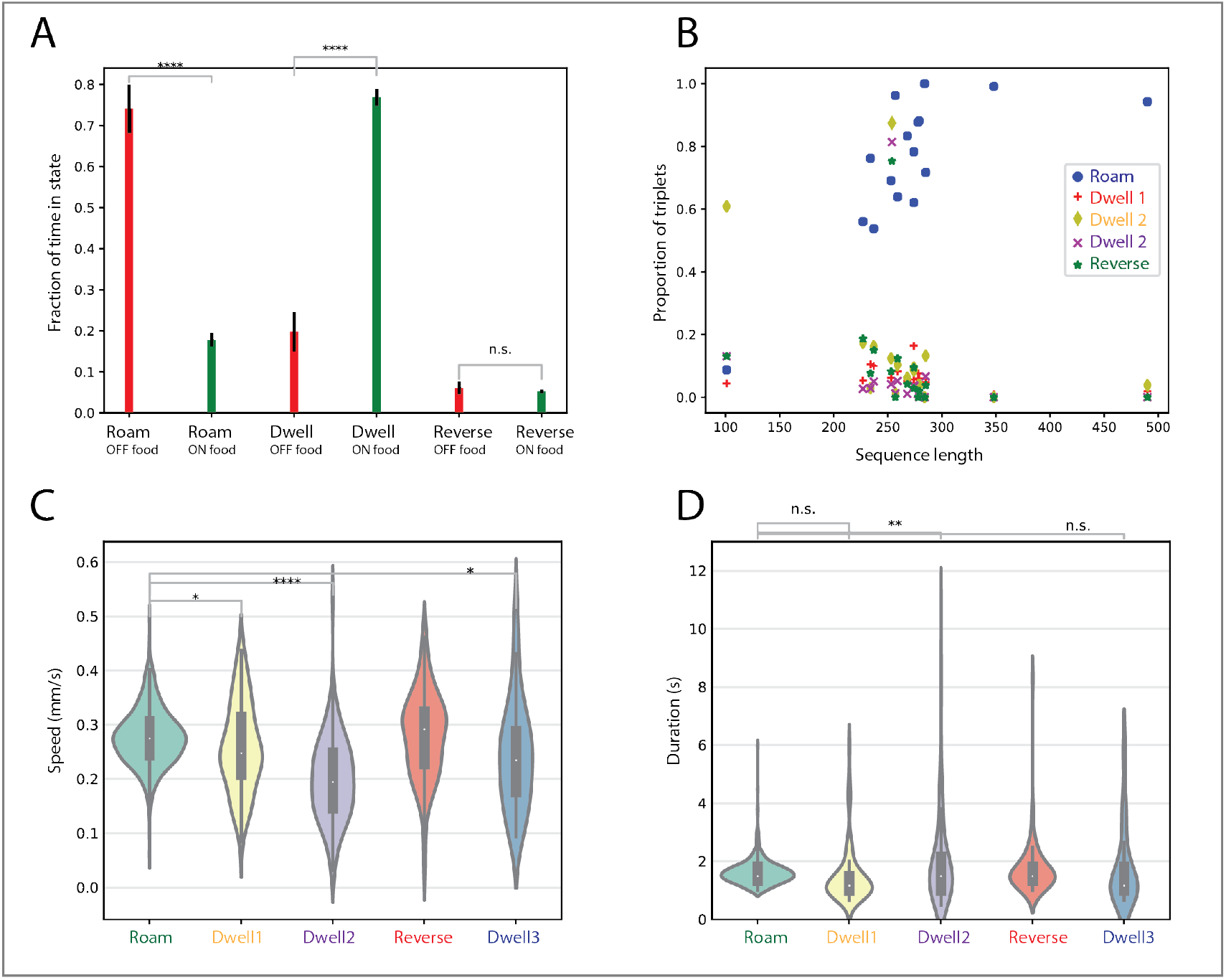
Grammatical rules underlying worm foraging capture behavioral variation induced by environmental changes. (**A**) Fraction of time spent in roaming-type and dwelling-type rules by N2 worms in off food (in red) versus on food (in green) conditions (mean ± s.e.m. across all the worms per condition; d(roam(N2off),roam(N2on)) = 3.56, d(dwell(N2off),dwell(N2on)) = −3.94; N(N2off) = 16, N(N2on) = 30). (**B**) The proportion of roaming-type rule increases as a function of postural sequence length in both off food and on food worms. (**C**) Average speed for each instantiation of rule type across all worms off food (d(roam,dwell1) = 0.36, d(roam,dwell2) = 1.44, d(roam,dwell3) = 0.64; N(roam) = 1217, N(dwell1) = 92, N(dwell2) = 120, N(reverse) = 81, N(dwell3) = 39). (**D**) Time to complete individual instantiations of each of the behavioral rules across all worms (d(roam,dwell2) = −0.71; N values same as in (C)). Violin Plots contain box plots that show the inter-quartile range (**** = p < 0.0001, *** = p < 0.001, ** = p < 0.01, * = p < 0.05, Welch’s t-test; effect size via Cohen’s d).

A complementary way to visualize the difference between the behavior of N2 worms “on food” versus “off food” is shown in **Figure 17B**. For all the different lengths of posture sequences corresponding to worm behavior off food, the proportion of the roaming type rules (“b1→…→g3”) always exceeds that of the dwelling type rules (except for one worm whose posture sequence length is very low, approximately 100). Note that this is in contrast to **Figure 14A** depicting the behavior of N2 worms on food, where the proportion of roaming type rules remains in the range of 0.2 even when the length of the posture sequence exceeds 1000. This contrast exhibits how change in environment leads to a complete change in behavior and highlights how the proposed worm foraging grammar can account for such effects of environment on worm behavior.

The average speed during roaming type rules across all worms, in off food conditions, is significantly greater than the speed during dwelling type rules (**Figure 17C**) but the total time spent in roaming type rules does not differ from the time spent in Dwell 1 and Dwell 3 type rules (**Figure 17D**). This suggests that during the dispersal (roaming) period, the worm reorients itself for a very short time to change its direction (hence the low speed and relatively low time spent in dwelling type rules), to again continue with its roaming behavior.

Taken together, these results demonstrate the robustness of the proposed worm behavioral grammar in its ability to recapitulate variations in worm behavior that are caused by changes in its environment.

### The grammar reveals hitherto uncharacterized role of npr-3 & npr-10 genes in foraging

We next asked if the discovered worm foraging grammar can give insights into the molecular mechanisms underlying foraging. Neuropeptides and their receptors have been shown previously to affect foraging behavior. *npr-1* (ad609) mutants show increased roaming behavior in the presence of food (**Reddy et al., 2011; Gloria-Soria and Azevedo, 2008; De Bono and Bargmann, 1998; Cheung et al., 2004**), while *npr-9* (tm1652) mutants show impaired roaming behavior on food (**Bendena et al., 2008**).

We first sought to confirm if our proposed worm grammar for roaming and dwelling can replicate these findings. As shown in **Figure 18A**, we found that the total time spent by *npr-1* mutants in roaming type grammatical rules is significantly higher than wild type N2 worms. Furthermore, the amount of time spent by *npr-9* mutants in roaming type rule is significantly lower than N2 worms (**Figure 18B**). Note that the time spent in roaming type rules is the time spent in all rules of type “b1→…→g3” in a worm whereas the time spent in dwelling type rules is computed as the combined time spent in Dwell 1, Dwell 2 and Dwell 3 type rules.

**Figure 18.**
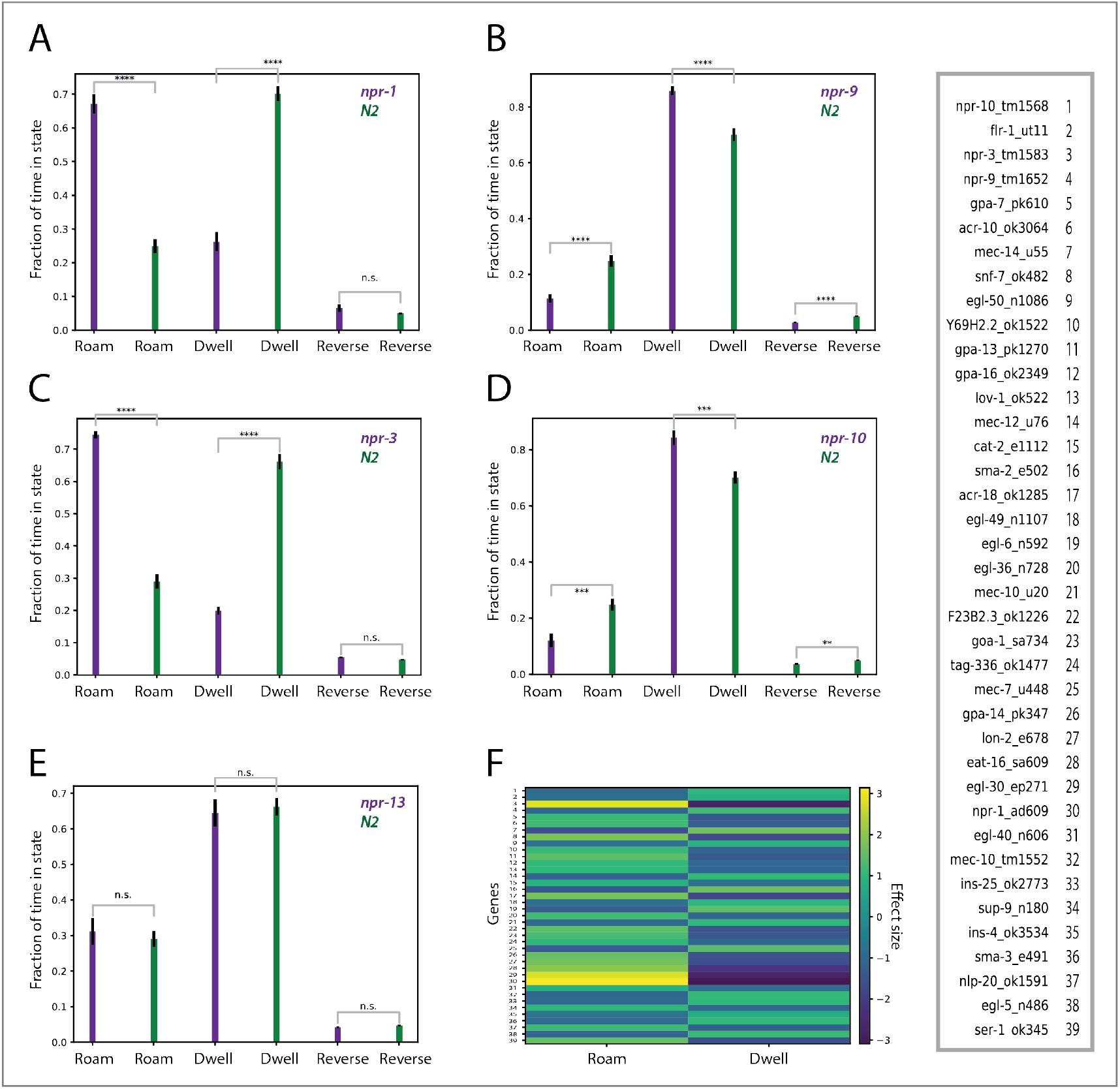
Time spent in roaming and dwelling grammatical rules reveals known and novel genes underlying worm foraging behavior. (**A**) Fraction of total time spent in roaming and dwelling type rules in *npr-1* mutants (d(roam(npr-1),roam(N2)) = 3.14, d(dwell(npr-1),dwell(N2)) = −3.08; N(npr-1)=12, N(N2)=44). (**B**) Fraction of total time spent in roaming and dwelling type rules in *npr-9* mutants (d(roam(npr-9),roam(N2)) = −1.1, d(dwell(npr-9),dwell(N2)) = 1.2; N(npr-9) = 28, N(N2) = 44). (**C**) *npr-3* mutants spend significantly higher time roaming as compared to N2 worms (d(roam(npr-3),roam(N2)) = 2.84, d(dwell(npr-3),dwell(N2)) = −2.79; N(npr-3) = 20, N(N2) = 61). (**D**) *npr-10* mutants spend significantly lower time roaming as compared to N2 worms (d(roam(npr-9),roam(N2)) = −0.95, d(dwell(npr-9),dwell(N2)) = 1.00; N(npr-10) = 21, N(N2) = 44). (**E**) *npr-13* mutants don’t differ in the time they spend in roaming and dwelling states as compared to N2 worms (*N(npr-13)=29, N(N2)=6).* All results shown as mean ± s.e.m., across all individual worms per strain. N values denote the number of worms considered for each mutant as well as N2 worms (**** = p < 0.0001, *** = p < 0.001, n.s. = not significant, Welch’s t-test; effect size: d = cohen’s d). (**F**) Other mutant strains screened that differ from N2 worms significantly in the time spent roaming and dwelling as given by the rules of worm foraging grammar (p value (Welch’s t-test) < 0.001 and |Cohen’s d| > 0.8). Colors in each indicate the actual effect size value (Cohen’s d) for the comparison between the time spent in either roaming or dwelling states for each of the mutant strains and N2 worms. Yellow values indicate the behavior type in which the time spent by the mutant strain is considerably higher as compared to N2 worms and vice versa for dark blue values.

For each worm, total amount of time spent in all roaming and dwelling type rules is used to calculate the proportion of total time spent roaming and dwelling. Building on these findings, we next sought to find if the worm grammar can help implicate hitherto uncharacterized genes that affect worm foraging behavior. We found that *npr-3* (tm1583) mutants show significantly increased propensity to roam (**Figure 18C**) whereas *npr-10* (tm1568) mutants show an opposite tendency to dwell (**Figure 18D**), as compared to N2 worms. It should be noted that not all neuropeptides and their receptors show significant differences in their foraging patterns as compared to N2 worms. For example, we found no significant difference in roaming and dwelling between npr-13 mutants and N2 worms (**Figure 18E**).

Finally, we looked at more than 300 worm strains and isolated those strains that differ significantly (p-value < 0.001, Welch’s t-test accompanied with a large effect size, Cohen’s d > 0.8) from N2 worms in the time spent in roaming and dwelling states as determined by the grammatical rules (**Figure 18F**). Yellow values (exact values equal to Cohen’s d statistic) indicate the behavior type in which the time spent by the corresponding mutant strain (on the y axis) is considerably higher as compared to N2 worms and vice versa for dark blue values. We can see that apart from the mutant strains belonging to the neuropeptide class elucidated above, *egl-30* mutants as well as mutants belonging to the *mec* class (*mec-14, mec-10, mec-12* and *mec-7*) show strongly different foraging patterns than N2 worms. Mutation in *egl-30* (ep271), known to be involved in chemosensory behavior and locomotion, leads to worms spending considerably more time in roaming than N2 worms. In turn, mutation in *mec-14* (u55), known to be involved in mechanosensory behavior, results in worms spending considerably more time in dwelling than N2 worms.

Prior work (**Flavell et al., 2013**) has also implicated *mod-1* gene in regulating roaming behavior in worms, however we did not find any significant difference in the foraging patterns between *mod-1* mutants and N2 worms, according to our proposed grammatical rules. We think that this could be because of the difference in experimental conditions, for example, the data we use involves a habituation period of 30 minutes before the actual worm tracking begins. This habituation period is absent in the study above. We must also note that the previous study (**Bendena et al., 2008**) that found that *npr-9* mutants have impaired roaming behavior, a result that we also find using our behavioral grammar, does involve a habituation period of 30 to 60 minutes, thus making their experimental conditions more similar to the experimental conditions of the behavioral data that we use.

These findings demonstrate that organization principles encapsulated by the proposed grammatical rules capture essential aspects of worm foraging in a way that can end up distinguishing between worm strains based on behavior. Moreover, the importance of delineating such organization principles is highlighted by their ability to elucidate novel molecular mechanisms important for regulating worm foraging behavior.

### The relational properties of behavioral rules remain invariant across mutant strains

We then looked at the properties of roaming and dwelling rules derived from N2 and wild isolates in a variety of available mutant strains. Following the division of various mutant strains into classes as described in (**Brown et al., 2013**), we found that the roaming and dwelling rules have qualitatively similar properties in all the mutant strains. Specifically, the relationship between the usage frequency of various kinds of rules with the behavioral sequence length follows the same pattern as in the wild isolates and N2 worms. Proportion of roaming type rules increases as a function of postural sequence length and vice versa for dwelling type rules (Dwell 2) for all the mutant strain classes analyzed (**Figure 19**).

**Figure 19.**
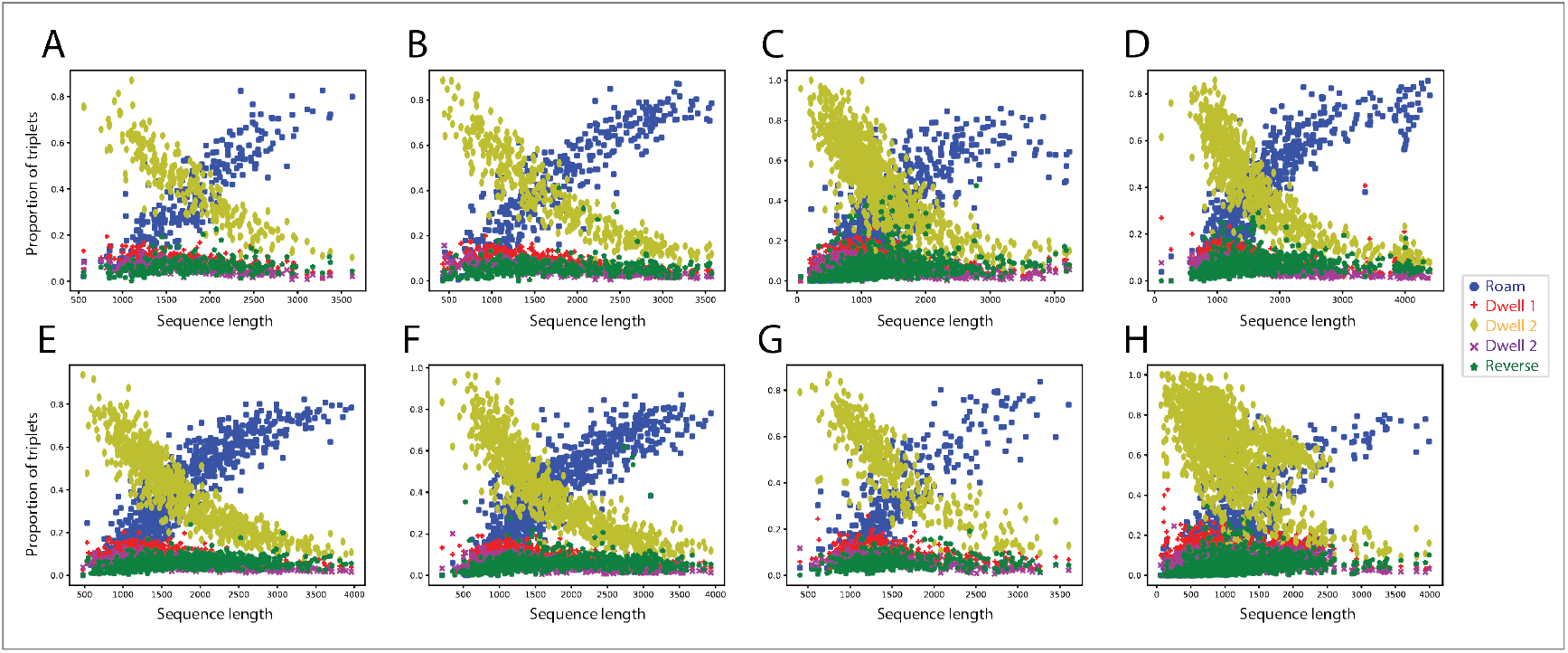
Similar usage frequency of roaming and dwelling rules across different classes of mutant strains. (**A**) Acetylcholine Receptor (N=268 worms). (**B**) *Deg/Enac* Channel (N=581). (**C**) *Egl* (N=969). (**D**) G-protein Related (N=741). (**E**) Monoamine Related (N=740). (**F**) Neuropeptide (N=881). (**G**) *Trpc* (N=373). (**H**) *Unc* (N=1412).

As with N2 and wild isolate worms, we found that the speed of the worm centroid for all the mutant strain classes during (**Figure S2**) the roaming type rules is significantly greater than that during dwelling type rules. These results suggest that the meaning of the rules (both roaming specific versus dwelling specific), despite quantitative differences, remains invariant across mutant strain classes; roaming specific rules having properties different from dwelling specific rules in precisely the manner as would be expected from the functional properties of roaming and dwelling worm behavior. Moreover, the total time spent in roaming type rules across all worms of a particular class is significantly smaller than that spent in dwelling rules (**Figure S3**).

### A context-free grammar (not a regular grammar) better captures worm foraging behavior

Overall, these results point to the existence of grammatical rules governing *Caenorhabditis elegans* foraging behavior, where a grammar specifies rules for transforming higher order chunks into a sequence of postures. For example, roaming behavior can be transformed into “B→R→G” and so on to generate posture sequences that lead to forward locomotion.

It is now appropriate to briefly pause here to mention an important notation distinction. An arrow between symbols (denoting modules or sub-modules) usually means a temporal transition rule. However, from a bottom-up syntactic view, elements get combined to form other constituent at higher levels. The arrow then signifies the celebrated “merge” operation. From a top-down view of syntax, the arrow implies command or contain. So, in the following technical specification of a grammar for the worm, the common use of arrows (transitions; A goes to B) will be in conflict with its syntatic use (going down the tree).

Thus, one way to specify transformation rules is, as shown below, where each symbol on the left of the symbol “→” generates (and so, is substituted) by the symbols on the right hand side of “→”. Symbols that never occur to the left of any grammatical rule (in other words, those symbols that cannot generate any new symbols) are known as terminal symbols. All the other symbols are known as non-terminal symbols. For example, the following two toy grammars can produce strings of type “(a^n)(b^n)”, where n are integer numbers:

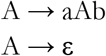

where A is a non-terminal symbol and a, b and ε (which denotes empty or null symbol) are the terminal symbols. A grammar generates multiple strings of terminal symbols, each of which is part of the language generated by the grammar. Thus, the strings, “ab”, “aabb”, “aaabbb”, are all part of the language generated by the two grammars above.

Keeping these preliminaries in mind and combining them with the knowledge of roaming and dwelling gleaned from the previous sections, we can formulate the following grammatical rules for *Caenorhabditis elegans* foraging, as recapitulated in **Figure 20**. Each of the 90 postures serves as a terminal symbol. All the other symbols are the non-terminal symbols. And ε = {} is an empty symbol.

**Figure 20.**
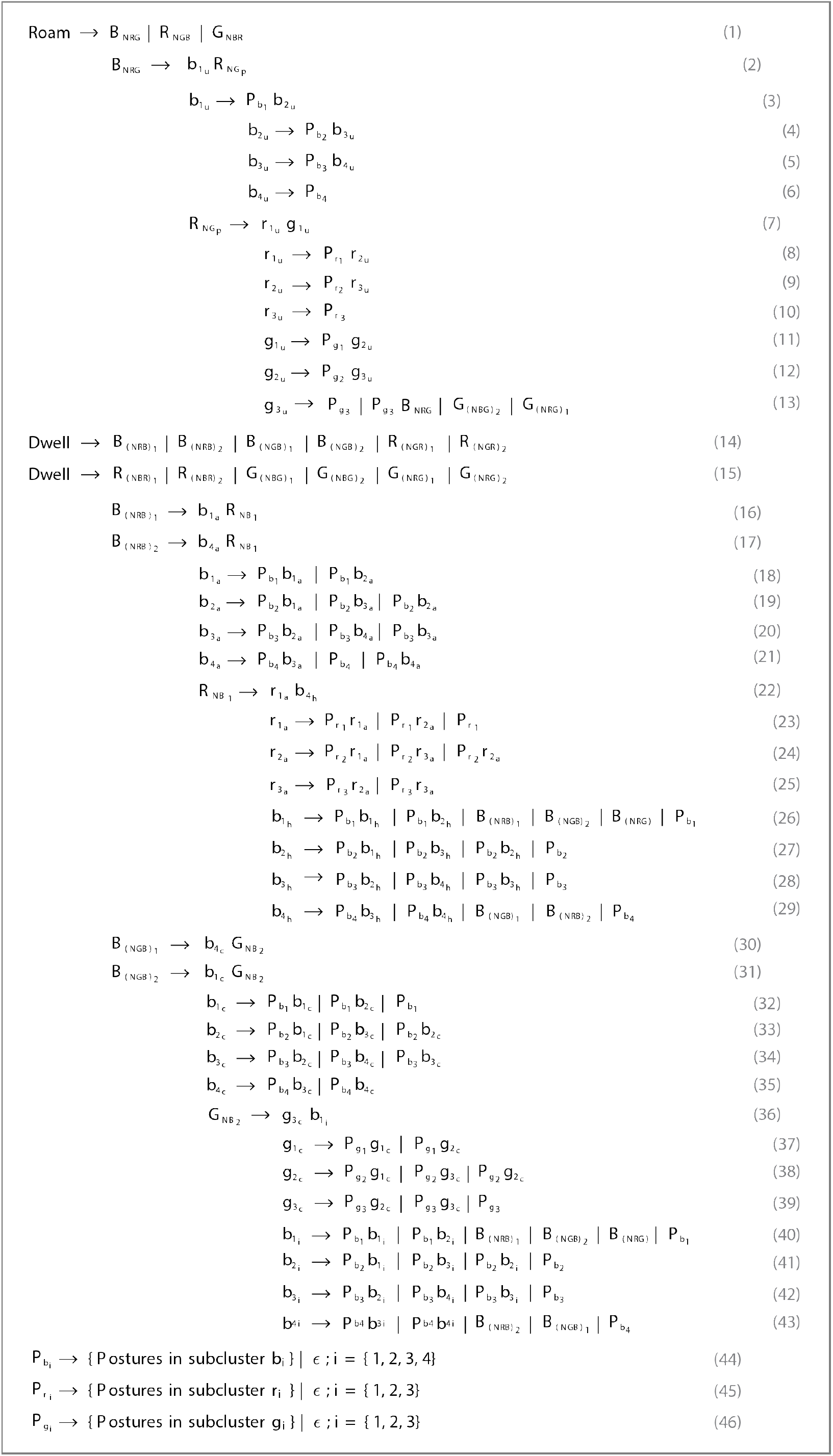
Grammatical rules for Caenorhabditis elegans behavior.

**Figure 21A** shows how the grammatical rules are instantiated to generate a posture sequence that implements worm roaming behavior (posture sequence {58, 18, 42, 24, 50, 19, 46, 10, 81, 26}). The symbol ‘|’ stands for the logical OR operator denoting choice in the expansion of non-terminal symbols in a grammatical rule. For example, **Equation 19** denotes that b_(2_(a)) can either be substituted by P_(b_(2)) followed by b_(1_(a)) or P_(b_(2)) followed by b_(3_(a)) or P_(b_(2)) followed by b_(2_(a)). **Figure 21A** then shows the instantiation of a roaming postural sequence through the B_NRG (“B→R→G”) grammatical rule, but similar grammatical rules can be made for RNGB (“R→G→B”) and G_NBR (“G→B→R”), all of which are known to generate roaming behavior. **Equation 13** also depicts that after completing one bout of forward locomotion through the b1→g3 rule, the worm can switch to dwelling by entering the G→R→G rule (either G_(NRG)1 or G_(NRG)1). Note that **Equation 14** denotes that one way to achieve dwelling behavior is through the behavioral unit B→R→B (B_(NRB)1 and B_(NRB)2). Once inside the unit, the grammatical rules that follow lay the groundwork for generating dwelling behavior through “B→R→B” unit, i.e. the rules corresponding to **Equations 16 to 46**.

**Figure 21.**
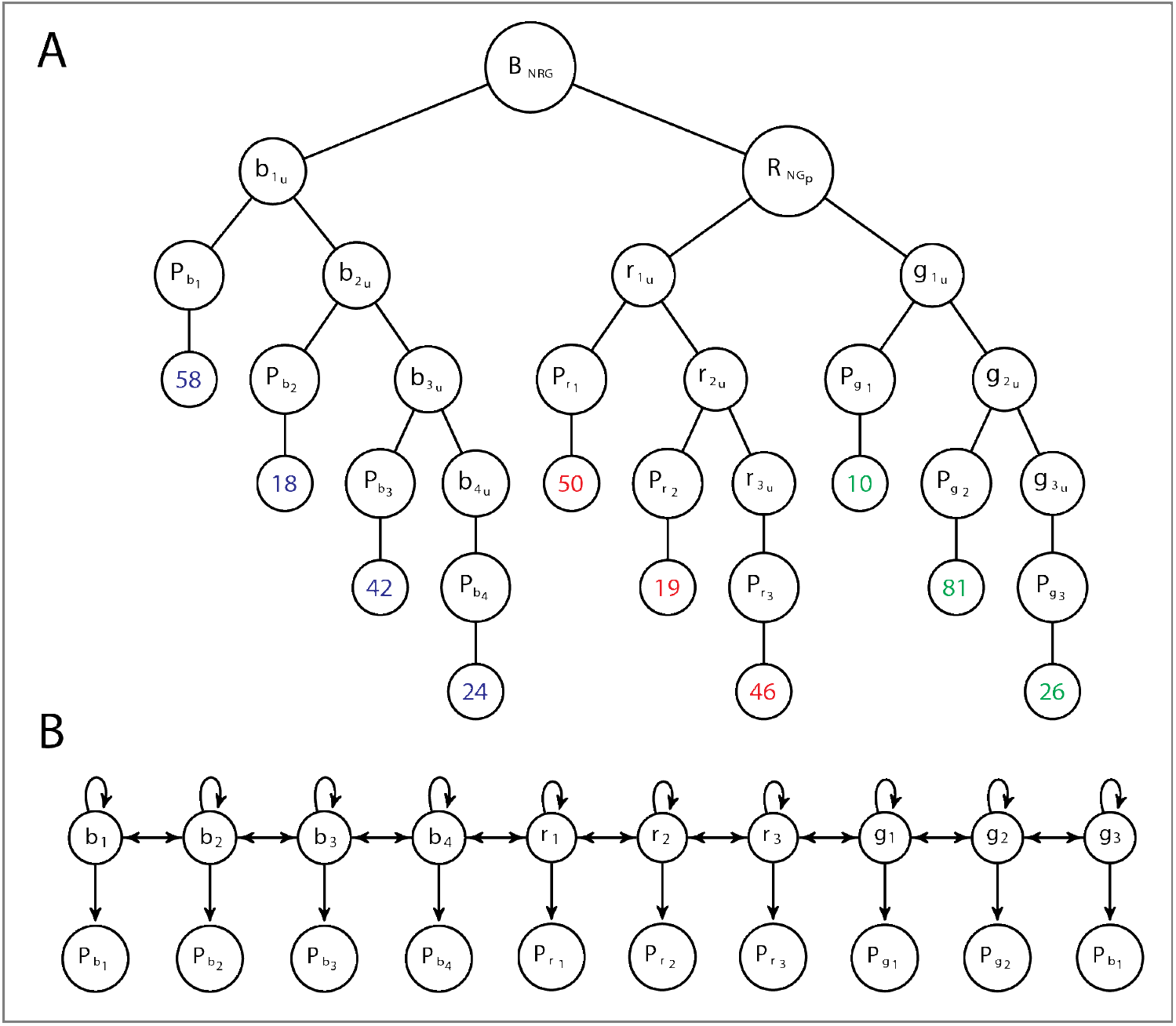
Conceptualizing behavior via context-free grammars versus hidden Markov chains. (**A**) Worm behavior can be generated by the application of specific grammatical rules. For instance, rules corresponding to Equations 1 to 13 in Figure 20 can generate the posture sequence “58, 18, 42, 24, 50, 19, 46, 10, 81, 26” that produces worm roaming behavior as shown in Figure 11D. Each of the postures in the sequence above serves as terminal symbols in the grammar, while the rest of symbols shown in the figure serve as non-terminal symbols. The grammar thus provides a way to transform non-terminal symbols into a sequence of terminal symbols according to set of production (grammatical) rules. (**B**) Hidden Markov Models (HMMs) approximate regular grammars by defining a linear relationship between hidden variables that account for actual observations, like worm postures. The transition rules between the hidden variables (in our case, the sub-modules b_i, r_i and g_i) remain the same, whether the worm is roaming or dwelling, which does not do justice to what is seen in worm behavior. When in roaming, the transition relationship between hidden variables should be different from the one used by the worm during dwelling, as exemplified by the worm itself and by the grammatical rules defined above.

The grammatical rules corresponding to other units like “R→G→R”, “R→B→R”, “G→B→G” and “G→R→G” can be generated in a similar manner. The assignment of individual postures to sub-clusters is done according to sub-cluster identities defined in **Figure 7C**. For example, P_(b_(1)) → {3, 20, 27, 41, 55, 58, 68, 76}, which means that whenever the pre-terminal symbol P_(b_(1)) is encountered, it is replaced by any of the postures (terminal symbols in our case) contained in the set of postures given by {3, 20, 27, 41, 55, 58, 68, 76}.

The rules described above constitute a context free grammar (**Chomsky, 1965**), whose underlying structure is hierarchical. This is because in a context free grammar, a non-terminal symbol on the left side can transform into (or generate) more than one non-terminal symbols (i.e. in rules given by **Equations 2** and **16**). And each non-terminal on the right hand side gets transformed into sequence of terminal symbols, rather than only a single terminal symbol.

In contrast, linear systems based on the Markovian assumption, for example, hidden Markov models (HMMs) approximate a regular grammar (**Chomsky, 1965**), where each non-terminal symbol on the left hand side of a rule does not get transformed into more than one non-terminal symbol on the right hand side. For example, if we were to consider a grammar for worm foraging that consisted only of the following grammatical rules, then we would have a regular grammar. Note that we have only specified the grammatical rules for B cluster and not shown the rules for the R and G cluster that can be made in a similar manner:

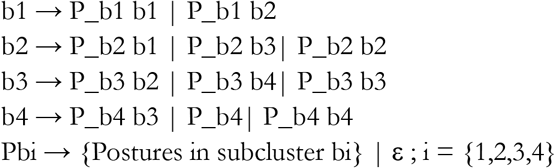

Note that P_b1, P_b2, P_b3 and P_b4 are not actually considered as non-terminal symbols, but rather pre-terminal symbols as they only generate terminal symbols (**Equation 44**) with no associated non-terminal symbols and are only ever used in the same way.

An HMM can be considered as a stochastic version of a regular grammar (the five rules in the previous paragraph) as shown in **Figure 21B**, where the non-terminal symbols b1, b2, b3 and b4 correspond to hidden states and each hidden state generates a posture with some associated probabilities and transitions to another hidden state. For example, the second rule in the last paragraph says that the worm in hidden state b2 generates any posture belonging to Pb2 (according to some probability distribution) and then the worm transitions to the hidden state b1 or b3 or stays in the same hidden state, based on certain probabilities.

Note that in this conception, there is no memory in the system, that is, there would be no difference in the transition probabilities between b1, b2 etc. depending on whether the worm is roaming or dwelling. However, we know that when the worm is roaming the probability of the worm transitioning from b2 to b1 is very close to 0 and the probability of transitioning from b2 to b3 is close to 1. And the worm follows different transition probabilities between the 10 sub-clusters when in dwelling phase. This is the problem with regular grammars and by extension HMMs, where they fail to account for structure that is hierarchical and only capture linear relationships.

On the other hand, a context free grammar gives a better account of worm behavior because the identity of b1 and hence the transition rules that it participates in, is intimately tied to the higher order chunk that it belongs to. For example if b1 is instantiated as b1a (**Equation 18**), we know that the worm is dwelling according to B(NRB)1 (B→R→B unit) rule (**Equation 16**) and the transitions between the sub-clusters is governed according to **Equations 16 to 29**. Whereas if b1 is instantiated as b1u (**Equation 3**), we know that the worm is in roaming state BNRG (B→R→G unit) (**Equation 2**) and hence the transitions are different and are governed by rules given by **Equations 2 to 13**. This gives a hierarchical conception of behavior where the transition between lower level elements (whether sub-clusters or individual postures) is not fixed (like in Markov models), but rather where the identity of the higher order chunk that the organism is in, governs the transition patterns (rules) between the chunks below (**Fentress and Stilwell, 1973**). Thus, a generative grammar provides a scaffold for worm behavior, indicating sequences that can and cannot be generated by the worm.

## 4. Discussion

While the scientist measures behavior sequentially, the organism generates it serially. Such distinctions matter (**Gomez-Marin, 2019**). Ethology’s most popular analysis tool and conceptual framework is arguably the ethogram: a catalogue of behaviors are linked by arrows depicting the probability that one leads to another. Yet, it seems unlikely that this is really how nervous systems control actions. Half a century ago, Simon mapped complexity to hierarchy, whereby organization is constructed via interconnected nested modules (**Simon, 1962**). Later, Dawkins proposed hierarchical organization as a general principle for behavior (**Dawkins, 1976**), addressing Lashley’s serial order problem (**Lashley, 1951**). Starting at the top of the hierarchy, animals make global decisions, progressively making narrower sub-decisions, until each act is instantiated (i.e. I decide to go to the lab on a sunny Sunday, then I decide to ride my motorbike to get there, and so I walk to the closet to get my helmet). Ascending from the bottom, behavioral units get grouped into “chunks” and further into “chunks of chunks”. Expanding on what the traditional ethogram can capture, knowledge of the chunk in which the previous action occurs, increases predictability of the next action.

Animal behavior is not only organized, it is also meaningful (for the animal, at least). Focusing on the precise trajectories traced by an organism is often necessary, but insufficient to make sense of what it is really doing. One may wish to enrich low level descriptions with intuitions about the behavioral “meaning” that operates on top of such “microscopic” descriptions. It is thus enticing —and daunting— to look at animal behavior through an “analysis of meaning” (**Clarke & Crossland, 1985**). In language, Ferdinand de Saussure made suggestions to analyze its structure and meaning in terms of two kinds of relations between the words that make up the language (**Harre, 1976**): (i) “syntagmatic relations”: rules that define how individual words should be strewn together to generate higher order structures like phrase and sentences; and (ii) “paradigmatic relations”: rules that define how different words can be associated with each other based on form or meaning by playing the same role in a sentence.

In this work, we used the ideas about syntagmatic and paradigmatic relations to analyze the structure of behavior in the nematode *Caenorhabditis elegans*. We uncovered rules that define how worm postures and higher order chunks of worm postures are combined to generate worm locomotor and foraging behavior. Grammatical rules specify how chunks of postures can be re-used in different contexts to produce different behaviors, roaming or dwelling. Such a generative grammar of worm foraging demonstrates how a compositional hierarchy can generate flexible behavior.

As already emphasized above, hierarchical organization has long been thought as a general principle of behavior (**Tinbergen, 1950, 1951; Dawkins, 1976**) but clear demonstrations of its existence during spontaneous behavior are conspicuous by their absence (**Brown & de Bivort, 2018**). A further missing piece in understanding behavior is how hierarchical organization aids in generating flexible animal behavior. Treating worm behavior as a sequence of body postural changes, we demonstrated that its foraging behavior —not just the stereotypical portions— is hierarchically organized. To that end, we used mutual replaceability (**Dawkins, 1976; Maurus & Pruscha, 1973**) to obtain chunks containing mutually substitutable worm postures (paradigmatic relations between postures), just like in the context of verbal behavior, the adjective chunk contains mutually substitutable words like “black” and “white”. The resulting chunks are then used to demonstrate how the worm can generate flexible behavior. Specifically, we elucidated a grammar of worm foraging outlining rules of interaction between such chunks containing substitutable postures (syntagmatic relations). We found that the stereotypical worm roaming behavior (sustained forward motion, with few turns and pauses) is captured by a specific grammatical rule involving specific chunks in a particular order. Let us remark that even such stereotypical behavior is characterized by variability at the lowest level of postures. A single rule specifying interactions between specific chunks can generate flexible roaming behavior where there is flexibility in the choice of postures used by the worm from each chunk.

Stereotypy in behavior patterns has been used effectively in the past to demonstrate hierarchical organization underlying various stereotypic behaviors in *Drosophila* (**Berman et al., 2016**). While stereotypy is indeed a central concept in behavior, relatively less stereotypic behavior patterns can constitute up to half of the behavior in some animals (**Berman, 2018**). In this work we have tried to filled this gap in the context of non-stereotypical *Caenorhabditis elegans* dwelling behavior by delineating grammatical rules that specify how the same chunks (as the ones used in stereotypical worm roaming behavior) are used in different combinations to produce non-stereotypical dwelling (involving termination of sustained forward motion) like behavior patterns. Such a generative grammar for worm foraging demonstrates how flexible behavior emerges from a hierarchically organized behavioral scaffolding.

We have also shown that the properties of our proposed behavioral grammar are consistent with known experimental results about the behavior of various mutant strains as well as the behavior of wild isolates that encounter changes in their environment. We report what we believe to be a hitherto uncharacterized role of neuropeptide receptors *npr-3* and *npr-10* in modulating *Caenorhabditis elegans* foraging behavior.

### Provisos of this work

Despite our attempts to build generative models that yield a deep understanding of animal behavior, our work has several limitations. In its current form, our proposal is not fully generative in its design. Our grammar does not capture richer statistical characteristics of behavior that can specify when a simulated worm should shift from roaming to dwelling. Also, our grammar does not have adequate mechanisms built-in to specify which of the multiple rules that form part of Dwell 2 type dwelling rules should be used at a particular instant to generate dwelling behavior. It seems that these aspects are statistical in nature. Such structure could in principle be captured by hidden Markov models (HMMs), as in (**Wiltschko et al., 2015**) or via recurrent neural network (RNNs) models (**Li et al., 2017**). Although immensely powerful in their ability to capture statistical regularities, HMMs do have the limitation of being linear and memory-constrained and hence can fail to capture longer time relationships in behavioral sequences (**Collado-Vides et al., 1996**). Although one can in principle learn hierarchical HMMs, the optimization underlying such learning process is liable to getting stuck in local minima, with considerable chances of learning the wrong structure (**Collado-Vides et al., 1996**). One avenue for future research would be to identify structure beforehand (like the grammar identified in this work) and then feed this structure to learning algorithms like HMMs or RNNs, such that both the inherent structure and the statistical regularities within such structure can be leveraged simultaneously to build powerful generative models of behavior.

In using the very convenient representation of worm behavior as discrete postural sequences developed in (**Schwarz et al., 2015**), we chose an “alphabet” of 90 template postures, also along the lines of previous work where the Minimum Description Length principle was used to extract hierarchical structure out of discrete postural sequences (**Gomez-Marin et al., 2016**). Such 90 template postures were found to capture more than 80% variance in the whole repertoire of postures actually taken by the worm (**Schwarz et al., 2015**). Although accounting for a sizable variance, the 90 templates still leave out some postures that the worm takes during foraging. To check whether our findings are qualitatively immune to such changes in the number of template postures used to capture worm locomotion, we changed the number of template postures from a small (N=45) to a high (N=150) “alphabet”. We found that our main results are qualitatively the same. Similar patterns characterize the transition matrix between the sub-modules (see Supplementary **Figure S1**).

To demonstrate that *Caenorhabditis elegans* foraging behavior is hierarchically organized, we performed clustering based on substitutability criteria to obtain chunks of postures. However, the fact that one can cluster behavior into chunks does not necessarily imply that the underlying organization is actually hierarchical. This limitation was consciously present in previous work of ours (**Gomez-Marin et al., 2016**), being one of the main conceptual spurs of the present work. Yet, this apparent limitation in the present work is mitigated by complementing the knowledge of behavioral chunks with a behavioral grammar that specifies how the chunks obtained by clustering might be used to actually generate behavior sequences by the worm. This conception of hierarchical organization is in line with Herbert Simon’s notion of hierarchy (**Simon, 1962**), in which he argued that hierarchy does not only mean obtaining partitions of data but it also involves elucidation of the rules of interaction between the obtained partitions.

Studying behavior at the level of body postures is convenient for several reasons. First, by anchoring the level of analysis at the confluence of neurons, muscles and the environment, it tames the temptation to disembody the animal and its behavior. Moreover, in the case of the worm, its particular body plan induces a tight correlation between egocentric and allocentric spaces: changes at the level of postures are translated nearly univocally into changes at the level of centroid trajectory. In other words, one can map posture-motion to loco-motion (**Stephens, 2008; Keaveny & Brown, 2017**). However, in this work we worked at the level of postures due to the handy discrete representation at hand, which allows us to treat “behavioral data as text” which, in turn, lends itself to import powerful concepts and methods from other sciences. We have not yet studied in depth the role of body morphology in foraging beyond showing that the postural shapes, by themselves, cannot explain the transitions observed. Of course to transition from one postural template to another, such templates need to be close-enough in morphological space. But an important question still begs: why do some postures belong to a sub-module, and not others? And, why are there postures so morphologically distinct within the same module? In future work one may try to establish a concrete analogy with language again, this time getting inspiration from phonology (i.e. in the final voicing of pairs such as “d” and “t”, “b” and “p”, or “g” and “k”, the elements that form the group are grouped together in a natural class where rules still apply if you replace them). One can also envision another analogy with acoustic analysis, since how a phoneme sounds is different than the articulation needed to produce it. In this sense, not only neurons but also the role of muscles —and their specific relation both to neurons and to changes in body posture— would provide further hints into the organization of worm behavior. *C. elegans* is undoubtedly a fantastic organism to study these matters (**Gjorgjieva et al., 2014**), where plenty is known regarding the genetic and physiological mechanisms enabling its locomotion. There has recently been progress in integrating neural, mechanical and environmental aspects of worm locomotion by means of concrete and elaborate models (**Izquierdo & Beer, 2016; Izquierdo & Beer, 2018**). This highlights the importance of a comprehensive view of behavior within models that generate, rather than just recapitulate, behavior (**Izquierdo, 2019**). In sum, embodied “models that behave” provide further insights than “models of behavior” in a vat (**Gomez-Marin, 2017**).

### Further insights

The notion of substitution has allowed us to marry flexibility (of behavioral generation) with hierarchy (of behavioral organization) in order to elucidate a behavioral grammar for the worm. The generative model afforded by the behavioral grammar proposed here takes into account different timescales at which animal behavior manifests itself. This contrasts to other powerful generative models of behavior that operate at a single time scale (**Wiltschko et al., 2015**).

Furthermore, earlier attempts at obtaining a behavioral grammar have been characterized by human defined action labels, which can hinder the reproducibility of the analysis due to the biases inherent in human classification. Here, we tried to reduce such limitation by working with a representation that is comparatively less biased (and also more “microscopic”), form which to naturally spell out behavioral rules. Such approach based on clustering of substitutable behavioral descriptions should be applicable across organisms in a relatively robust manner. When applied to humans, it could provide a fairly precise dynamical understanding of behavior, revealing subtle changes in its organization upon aging and disease.

Whilst stereotypy may be considered as a general principle of behavior (**Berman et al., 2014**), a great deal of animal behavior consists of engaging in non-stereotyped behaviors. Thus, even if stereotyped behavior is shown to be hierarchically organized, one still needs to account for the organization of its non-stereotyped components. Here we have demonstrated that all the portions of *Caenorhabditis elegans* foraging behavior (both roaming, which is more stereotyped, and also dwelling, which is less stereotyped) are hierarchically organized.

It is often an expressed lament that, in spite of extensive knowledge about the anatomy and connectivity patterns in the nervous system of the worm, we still have not found ways to predict its behavior with sufficient accuracy at a sufficient level of detail. When studying animal behavior —which is modulated by a variety of factors, including the environment and the animal’s internal state—, one has to choose the level of abstraction at which prediction is to be sought. This becomes even more pertinent in scenarios where we study unrestrained behavior for a relatively long time, rather than a situation systematically paired with the so-called stimulus. In such unrestrained situations, the nature of predictions about behavior takes a different form than, let us say, “stimulus-response” behavioral paradigms (**Hofstadter, 1996; Hayek, 1964**).

Hierarchical organization helps in taming complexity by providing predictability at a higher level of abstraction, as opposed to lower levels of representation (postural templates, in our case) (**Fentress & Stilwell, 1973**). It follows that, in a hierarchically organized behavior, the predictability about the identity of lower levels of representation increases if we also know the chunk in which the behavior is currently happening. For example, if we know the worm is engaged in forward locomotion as in roaming states, we can deduce that it is in the B→R→G regime. If we further know it is roaming and currently in the B module, then we can deduce that it is in either of sub-modules b1, b2, b3 or b4, with the particular ordering of b1→b2→b3→b4. Finally, if we also know that the roaming worm is in the b1 sub-module, then our prediction ability improves further, because we know that there are only a few postures from which one choose, reducing the search space from the 90 original postures. In complement, flexibility is afforded via the relative freedom in the identity of posture that is used from each sub-module, since they are obtained by looking for substitutable postures in the first place (type 2 substitution). Another way in which predictability and flexibility can coexist comes in the system is through the re-usability of the same structures in generating different behaviors like roaming and dwelling (type 1 substitution).

### Future directions

One cannot avoid pondering over the neural dynamics subservient to hierarchically generated flexible behavioral dynamics. The concepts of degeneracy and re-usability that we have leveraged to tie hierarchical organization with flexible behavior also apply at the interface between the nervous system and behavior in the worm. For example, octanol-avoidance behavior is driven primarily by ASH nociceptive neurons in well-fed worms. But after an hour of starvation, the same behavior is mediated by ASH, AWB and ADL nociceptive neurons (degeneracy) (**Chao et al., 2004**). Moreover, differential activation of *npr-1* neuropeptide receptor allows ASH neurons to generate different behaviors by facilitating two different neuromodulatory states, with aerotaxis behavior occurring irrespective of modulation and aggregation behavior occurring only when *npr-1* activity is low (re-usability of the same neural circuitry to generate different behaviors) (**Cheung et al., 2005; Chang et al., 2006**). It remains to be seen if degeneracy (type 2 substitution) and re-usability (type 1 substitution) can help elucidate degenerate and re-usable neural circuit mechanisms related to flexible behavior.

Studies investigating the relationship of neural activity with behavior in *Caenorhabditis elegans* have found that the whole brain dynamics lie on a low-dimensional manifold. Trajectories of neural activity through this manifold can then be mapped to different behavioral states in the worm (**Kato et al., 2015**). It has been reported that reasonably differentiated trajectories though this manifold can correspond to the same behavioral state like reversals. Stereotyped behaviors like roaming that are generated by a single grammatical rule with flexibility in the choice of postures might be expected to trace trajectories relatively close to each other. On the other hand, less stereotyped dwelling behavior is characterized by multiple grammatical rules. Thus, one could hypothesize that different grammatical rules —corresponding to Dwell 1, Dwell 2 and Dwell 3— should correspond to differentiated trajectories through the neural manifold.

Finally, our finding that *npr-3* and *npr-10* modulate worm foraging could be used to further investigate the neural dynamics underlying roaming and dwelling (**Gray et al., 2005**). Gaining knowledge as to how neuropeptides like *npr-3* and *npr-10* influence the known neural circuitry associated with roaming and dwelling could then provide a more complete understanding of how the concerted action of genes and neural circuits enables behavior.

Syntax has also an important role to play in understanding the evolution of behavioral flexibility. The ideas of hierarchical organization and a grammar have fertilized inquiries both in linguistics as well as ethology (**Peters, 1981; Lashley, 1951; Chomsky, 1965; Kalmus, 1969; Fentress & Stilwell, 1973**). Grammar —either in the context of motor behavior or linguistics— involves defining ways in which elements may be permuted or combined at different levels of hierarchical organization to generate different sequences flexibly. Due to such multi-level permutations, even a small increase in number of elements and rules of combination can lead to disproportionate increase in the number of novel behavioral repertoires. Suggestions have been made in the past about how behavioral flexibility in higher animals can be seen as a series of evolutionary steps where the fixed action patterns of simpler organisms are first isolated into component primitive units that are then recombined in a variety of ways (using rules of grammar) in descendant and more complex animals (**Peters, 1981**). Consequently, a similarity in the rules of combination of behavior units (grammar) between animals, irrespective of the identity of the behavioral units might indicate evolutionary relationship. This would drive researchers to look not only for homologues of behavioral elements but also of rules for combining behavioral elements (**Peters, 1981**).

### Making the “behavioral language” analogy concrete, but not literal

The fundamental question that guided this work has been how to account for the production of serial behavior beyond sequential frameworks. We have proposed the use of grammatical rules as a conceptual and methodological alternative to Markov chains (hidden or not); one that we believe makes more justice both to the data and to the nature of behavior from the organism’s point of view. And yet, one should not conflate models of reality with reality. The grammars proposed here can describe the behavior of the worm —and in a sense also generate it, while capturing essential properties— but they do not prescribe it *sensu stricto*. Nor is our attempt to link worm behavior to linguistics a proposition for a nematode language as it is meant in humans. Our work is unavoidably influenced by Chomsky’s work, and yet we are not putting forth “a Chomskyan worm”. One should be extremely prudent in ascribing a grammatical capacity to animals (be it a worm, be it a marmoset) in the sense it is meant in humans. Here we made the analogy concrete, not literal.

To end, let us suggest that nervous systems may have evolved to be able to produce behavioral grammars of the kind we found in the worm. And yet, despite our speculations above, when it comes to the physical (neural or otherwise) implementation of behavioral grammars, we suspend judgement. What is the ontology of these rules? We do not commit to the same ontological status that syntacticians do. Linguists usually think of modules in the brain responsible for such rules. If specific, where are they? If general, where do they come form? We have no answer for the worm. The question may even be ill-posed. Nor do we obligate an information principle. Be it as it may, it is tempting to think that the organization of behavior we found in the humble nematode could reflect a structure deeply ingrained in biology. Back to the insights of classical ethology, it seems that everything human has its roots in “lower” animal behavior.

## Supplementary material

All codes and data used in this study can be found at: https://github.com/bksgupta/c_elegans_behavioral_grammar

## Contributions

AGM conceived the project and acquired the funding; SG analyzed the data and developed the grammars; SG made the plots; AGM made the figures; SG wrote the manuscript; AGM reviewed and edited the final version of the manuscript.

## Acknowledgements

We thank André Brown, Alfonso Pérez-Escudero, Ibrahim Tastekin, Pedro Tiago Martins, Aitor Landete and Shruthi Ravindranath for feedback on the manuscript.

## Supplementary figures

**Figure S1.**
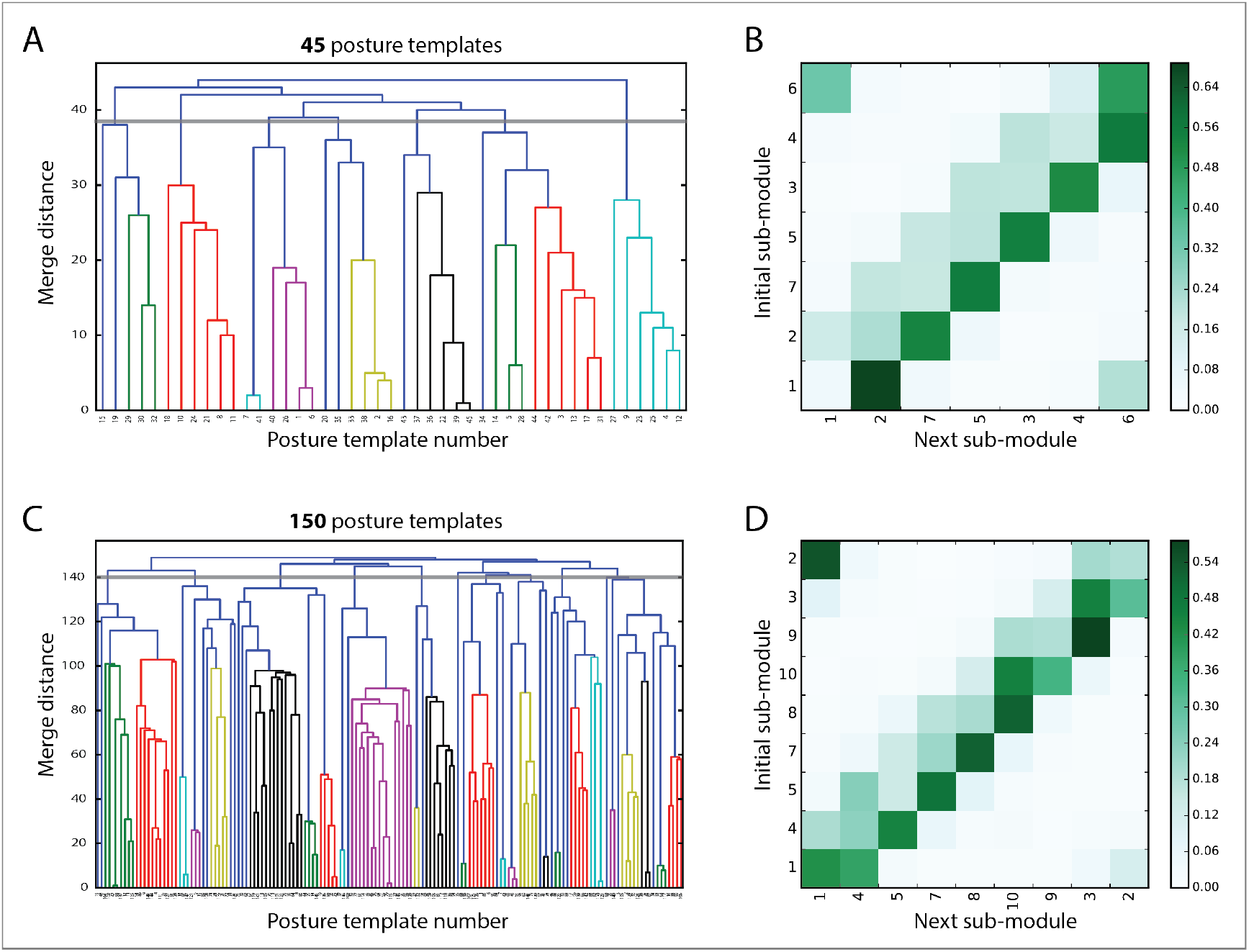
Results unchanged when choosing a larger or smaller set of postural templates. Decreasing or increasing the number of template postures used in the analysis of worm foraging behavior reveals the same patterns as seen with 90 postures. (**A**) Mutual replaceability dendrogram sub-module structure for N2 worms described with 45 posture templates. (**B**) Sub-module transitions for all N2 worms pooled together. (**C**) Dendrogram when using 150 templates. (**D**) Corresponding sub-module transitions.

**Figure S2.**
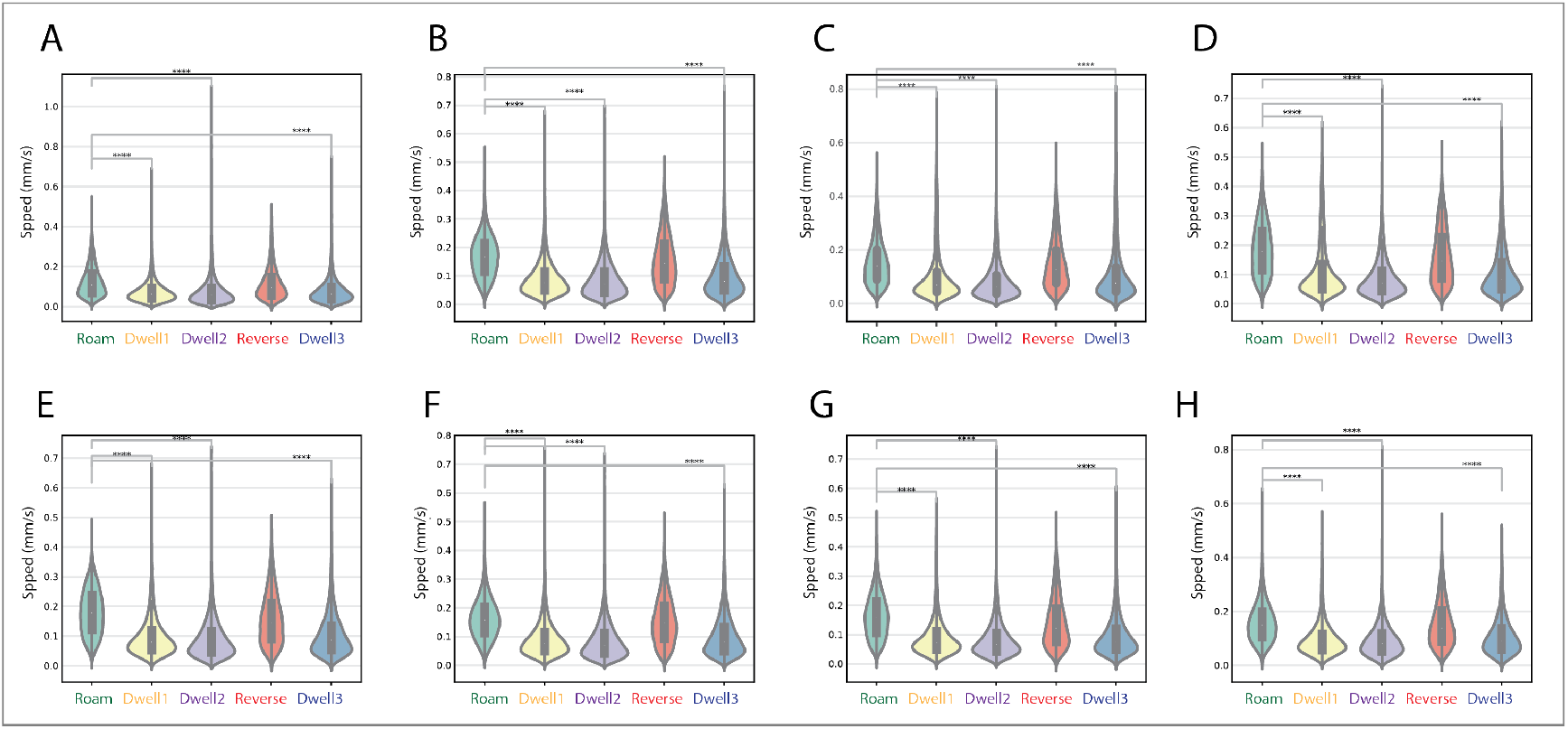
Average speed during instantiations of the roaming-type rule is considerably higher than during the behavior rules corresponding to dwelling for all the mutant strain classes: (**A**) Acetylcholine Receptor (268 worms); (**B**) *Deg/Enac* Channel (581 worms); (**C**) *Egl* (969 worms); (**D**) G-protein Related (741 worms); (**E**) Monoamine Related (740 worms); (**F**) Neuropeptide (881 worms); (**G**) *Trpc* (373 worms); (**H**) *Unc* (1412 worms). Violin plots showing the distribution of average speeds across all the worms. Box plots within show the inter-quartile range (**** = p<0.0001, Welch’s t-test; effect size: Cohen’s d). (A) d(roam,dwell1)=0.94, d(roam,dwell2)=1.02, d(roam,dwell3)=0.72. N(roam)=62178, N(dwell1)=12898, N(dwell2)=53980, N(reverse)=10122, N(dwell3)=8242. (B) d(roam,dwell1)=1.13, d(roam,dwell2)=1.22, d(roam,dwell3)=0.97. N(roam)=154215, N(dwell1)=22534, N(dwell2)=94293, N(reverse)=17699, N(dwell3)=12847. (C) d(roam,dwell1)=0.58, d(roam,dwell2)=0.939, d(roam,dwell3)=0.529. N(roam)=145805, N(dwell1)=40371, N(dwell2)=181788, N(reverse)=30975, N(dwell3)=24929. (D) d(roam,dwell1)=0.85, d(roam,dwell2)=1.25, d(roam,dwell3)=0.88. N(roam)=177213, N(dwell1)=33080, N(dwell2)=132831, N(reverse)=28067, N(dwell3)=19713. (E) d(roam,dwell1)=1.15, d(roam,dwell2)=1.34, d(roam,dwell3)=1.05. N(roam)=194924, N(dwell1)=32882, N(dwell2)=139543, N(reverse)=25179, N(dwell3)=20196. (F) d(roam,dwell1)=1.08, d(roam,dwell2)=1.2, d(roam,dwell3)=0.92. N(roam)=216351, N(dwell1)=36003, N(dwell2)=157452, N(reverse)=29545, N(dwell3)=21268. (G) d(roam,dwell1)=0.995, d(roam,dwell2)=1.23, d(roam,dwell3)=0.96. N(roam)=70560, N(dwell1)=17428, N(dwell2)=73622, N(reverse)=10557, N(dwell3)=9873. (H) d(roam,dwell1)=0.76, d(roam,dwell2)=0.8, d(roam,dwell3)=0.69. N(roam)=109010, N(dwell1)=58221, N(dwell2)=295363, N(reverse)=27357, N(dwell3)=33939.

**Figure S3.**
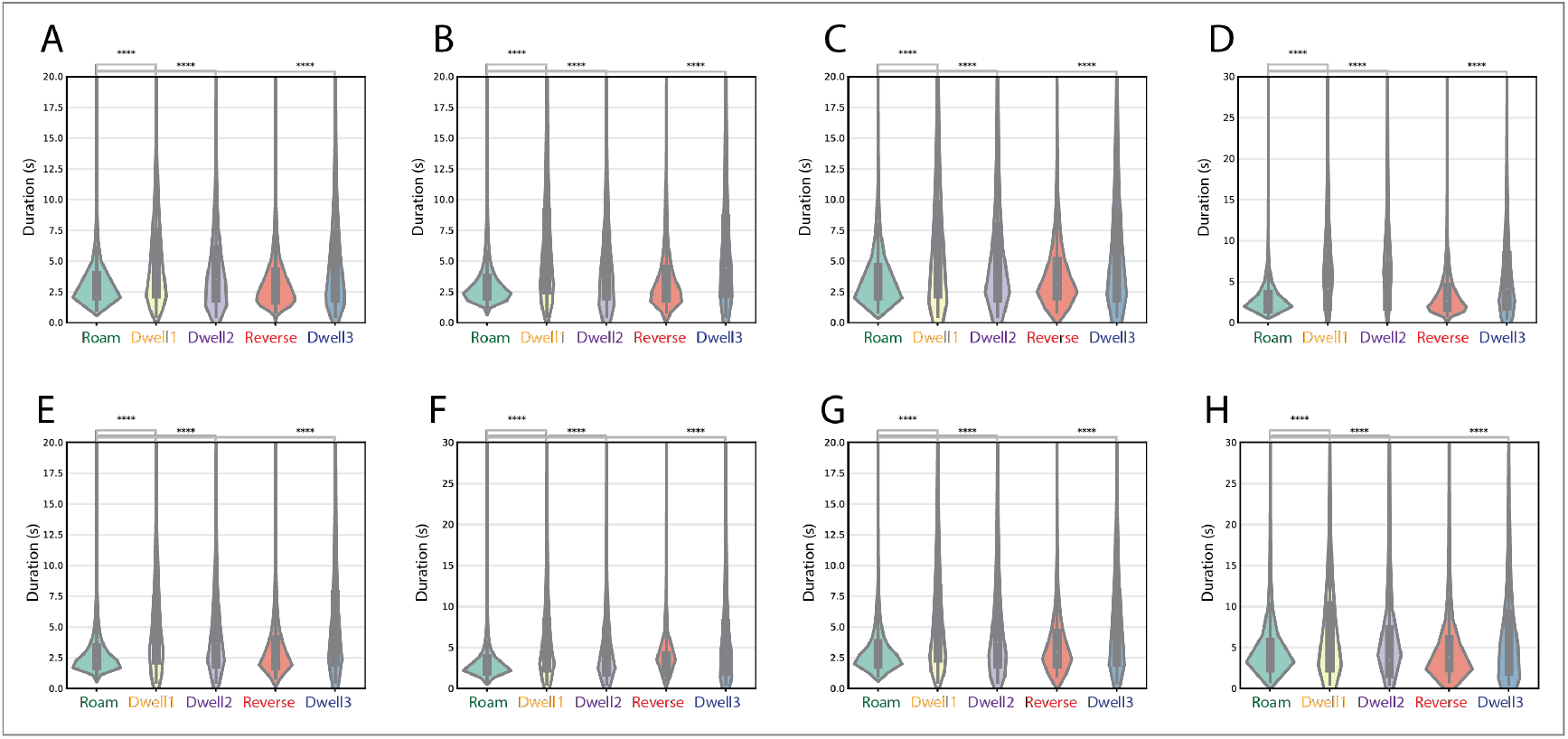
Roaming-type rules have a shorter time duration than dwelling-type rules for all the mutant strain classes: (**A**) Acetylcholine Receptor (268 worms); (**B**) *Deg/Enac* Channel (581 worms); (**C**) *Egl* (969 worms); (**D**) G-protein Related (741 worms); (**E**) Monoamine Related (740 worms); (**F**) Neuropeptide (881 worms); (**G**) *Trpc* (373 worms); (**H**) *Unc* (1412 worms). Violin plots show the distribution of times taken to complete each instantiation across all worms. Box plots show the inter-quartile range (**** = p<0.0001, Welch’s t-test; effect size: Cohen’s d). (**A**) d(roam,dwell1)=−0.89, d(roam,dwell2)=−0.43, d(roam,dwell3)=−0.86. (**B**) d(roam,dwell1)=−1.21, d(roam,dwell2)=−0.56, d(roam,dwell3)=−1.21. (**C**) d(roam,dwell1)=−0.74, d(roam,dwell2)=−0.39, d(roam,dwell3)=−0.78. (**D**) d(roam,dwell1)=−0.81, d(roam,dwell2)=−0.568, d(roam,dwell3)=−1.14. (**E**) d(roam,dwell1)=−1.09, d(roam,dwell2)=−0.6, d(roam,dwell3)=−1.26. (**F**) d(roam,dwell1)=−1.05, d(roam,dwell2)=−0.528, d(roam,dwell3)=−1.09. (**G**) d(roam,dwell1)=−0.97, d(roam,dwell2)=−0.54, d(roam,dwell3)=−1.02. (**H**) d(roam,dwell1)=−0.42, d(roam,dwell2)=−0.18, d(roam,dwell3)=−0.46.

